# Polarized localization of kinesin-1 and RIC-7 drives axonal mitochondria anterograde transport

**DOI:** 10.1101/2023.07.12.548706

**Authors:** Youjun Wu, Chen Ding, Alexis Weinreb, Laura Manning, Grace Swaim, Shaul Yogev, Daniel A. Colón-Ramos, Marc Hammarlund

## Abstract

Mitochondria transport is crucial for mitochondria distribution in axons and is mediated by kinesin-1-based anterograde and dynein-based retrograde motor complexes. While Miro and Milton/TRAK were identified as key adaptors between mitochondria and kinesin-1, recent studies suggest the presence of additional mechanisms. In *C. elegans*, *ric-7* is the only single gene described so far, other than kinesin-1, that is absolutely required for axonal mitochondria localization. Using CRISPR engineering in *C. elegans*, we find that Miro is important but is not essential for anterograde traffic, whereas it is required for retrograde traffic. Both the endogenous RIC-7 and kinesin-1 act at the leading end to transport mitochondria anterogradely. RIC-7 recruitment to mitochondria requires its N-terminal domain and partially relies on MIRO-1, whereas RIC-7 accumulation at the leading end depends on its disordered region, kinesin-1 and metaxin2. We conclude that polarized transport complexes containing kinesin-1 and RIC-7 form at the leading edge of mitochondria, and that these complexes are required for anterograde axonal transport.

**Summary statement:** Anterograde transport of axonal mitochondria is critical for maintenance of the mitochondria pool and neuronal health. Wu et al. show that the endogenous kinesin-1 and RIC-7 localize at the leading end of mitochondria to drive axonal anterograde transport *in vivo*.

## Introduction

Mitochondria are complex organelles with multiple functions including ATP production, calcium buffering, reactive oxygen species signaling, and programmed cell death. In neurons, synaptic mitochondria support neurotransmission by at least two mechanisms: serving as an energy source via ATP production, and acting as a reservoir for calcium buffering (Ivannikov et al., 2013; Kang et al., 2008; Kwon et al., 2016; Sun et al., 2013; Tang and Zucker, 1997; Verstreken et al., 2005). Therefore, maintaining mitochondria distribution is critical for neuronal homeostasis. Indeed, mitochondria mislocalization can lead to neurodegeneration, and defects in mitochondria trafficking are found in a variety of disease models for Alzheimer’s disease, Parkinson’s disease, and amyotrophic lateral sclerosis (Baldwin et al., 2016; De Vos et al., 2007; Ding et al., 2022; Lopez-Domenech et al., 2016; Nguyen et al., 2014; Rawson et al., 2014; Rui et al., 2006; Wang et al., 2010; Wang et al., 2011; Zhao et al., 2010).

Although evidence suggests that mitochondria biogenesis can occur in the axon, it is believed that most mitochondria are generated in the cell body and transported to distal processes (Amiri and Hollenbeck, 2008; Davis and Clayton, 1996). Mitochondria transport is primarily driven by the microtubule plus-end directed motor kinesin-1 and the minus-end directed motor dynein (Hurd and Saxton, 1996; Pilling et al., 2006). Microtubules in the axon are predominantly oriented with their plus end toward the distal axon. Therefore, kinesin-1 mainly drives the exit of mitochondria from the cell body into the axon and anterograde trafficking toward the distal axon, whereas dynein moves mitochondria retrogradely in the axon toward the cell body.

The kinesin-1 and dynein motors do not bind mitochondria directly, but through adaptor complexes. Miro and Milton/TRAK are key adaptors that link kinesin-1 and dynein to mitochondria (Amiri and Hollenbeck, 2008; Glater et al., 2006; Guo et al., 2005; Stowers et al., 2002; van Spronsen et al., 2013). Miro is a conserved GTPase with a C-terminal transmembrane domain that inserts into the outer mitochondria membrane (Fransson et al., 2003). Miro binds to Milton, a coiled coil protein that in turn binds to kinesin-1 and to dynein (Glater et al., 2006; van Spronsen et al., 2013). *In vitro*, Miro can also bind to KIF5/kinesin-1 directly (Macaskill et al., 2009). In Drosophila photoreceptors and motor neurons, loss-of-function of either Miro or Milton leads to dramatic loss of mitochondria in axons and depletion of mitochondria at the neuromuscular junctions, suggesting that Miro and Milton are essential for mitochondria exit from the cell body and anterograde trafficking in the axon (Guo et al., 2005; Stowers et al., 2002). By contrast, cultured hippocampal neurons from Miro knockout mice show normal amounts of mitochondria in the axon, with disturbed mitochondria distribution in the dendrite (Lopez-Domenech et al., 2016), suggesting that there are Miro-independent mechanisms to transport mitochondria into the axon in mammals. Indeed, a recent study suggests that Miro proteins are not required for TRAK/kinesin-dependent anterograde movement in mouse embryonic fibroblasts (Lopez-Domenech et al., 2018). These conflicting data suggest that our understanding of mitochondria transport *in vivo* is incomplete.

Studies in *C. elegans* also support the existence of Miro/TRAK-independent mechanisms of mitochondria transport. Although Miro and TRAK-1 are essential for dynein-based trafficking, kinesin-1-mediated transport doesn’t require them (Zhao et al., 2021). Miro is not required for axonal mitochondria distribution in some unipolar neurons such as touch receptor neurons and PVQ neurons (Ding et al., 2022; Sure et al., 2018). A recent genetic screen identified the metaxins MTX-1 and MTX-2, components of the mitochondrial sorting and assembly machinery, as important regulators for mitochondria transport (Zhao et al., 2021). In the DA9 neuron, genetic analysis indicated that *mtx-1/2* and *miro-1* redundantly regulate kinesin-1 mediated mitochondria trafficking, and that overexpression of *mtx-1 or mtx-2* in the PVD neuron can substitute for MIRO-1 for mitochondria transport by kinesin-1 (Zhao et al., 2021).

Another protein, RIC-7, is essential for mitochondria localization in *C. elegans*. In *ric-7* mutants, mitochondria are absent from the axons, similar to kinesin-1 mutants (Rawson et al., 2014). RIC-7 does not contain known conserved domains or motifs and does not have an obvious homolog in vertebrates (Rawson et al., 2014). RIC-7 is localized on mitochondria, but how it regulates mitochondria localization in neurons is not understood (Rawson et al., 2014).

Here, we analyze mitochondria transport *in vivo*, using the *C. elegans* AIY and DA9 neurons as model systems. We interrogate the mitochondria transport machinery by tagging the endogenous proteins and visualizing their localization and dynamics in physiological conditions, as well as by genetic analysis using deletion mutants. We find that Miro is important for robust mitochondria anterograde trafficking in axons but is not absolutely required, whereas it is essential for retrograde axonal trafficking. We demonstrate that RIC-7 is specifically required for kinesin-1-based mitochondria trafficking by localizing with kinesin-1 to the leading end of mitochondria. Further, Miro/MIRO-1 and metaxin2/MTX-2 function together to regulate the localization of RIC-7. Using structure/function analysis, we identify the domains in the RIC-7 protein that are crucial for its localization and function. Our work provides new insights into the cellular mechanisms that govern mitochondria traffic in living animals, and identifies polarized protein localization at the leading end as a novel functional site critical for organelle movement.

## Results

To assess mitochondria distribution *in vivo*, we validated a matrix-targeted EGFP marker by comparing the distribution of fluorescently tagged mitochondria in the AIY neuron with AIY mitochondria reconstructed from serial EM sections (White et al., 1986). The AIY neurons are a pair of unipolar interneurons that each extends a single neurite anteriorly, with microtubule plus ends orienting toward the distal end of the neurite (Hill et al., 2019), and have been well established as a model system to study presynaptic assembly and synaptic vesicle localization (Colon-Ramos et al., 2007; Stavoe and Colon-Ramos, 2012; Stavoe et al., 2012; White et al., 1986). We found that mitochondria distribution and overall morphology are similar between the EM data and our fluorescent label, indicating that the fluorescent marker accurately labels mitochondria without perturbing their native distribution *in vivo* (Fig. S1 and Methods).

### Axonal Anterograde Transport Does Not Require Miro or TRAK

In AIY, partial loss-of-function of *unc-116,* the sole ortholog of kinesin-1 *in C. elegans*, results in dramatic loss of mitochondria in the neurite (Fig. 1), consistent with an essential role for kinesin-1 in mitochondria anterograde transport in the axon (Hurd and Saxton, 1996; Pilling et al., 2006; Rawson et al., 2014). To assess the role of TRAK/Milton and Miro, we used CRISPR to generate deletion alleles of *trak-1,* the sole *C. elegans* TRAK ortholog, and of *miro-1, -2,* and *-3,* the three *C. elegans* Miro orthologs. We found that mitochondria localization is normal in AIY neurites in *trak-1(KO)* mutants and in a previously published *trak-1 loss-of-function* allele (Zhao et al., 2021) (Fig. 1), consistent with the finding that mitochondria distribution is not affected in the axon of the DA9 neuron in *trak-1* mutants (Zhao et al., 2021). Surprisingly, we found significantly more mitochondria in the AIY neurites of *miro-1(KO)* animals. This increase in the number of axonal mitochondria is not due to compensation from the other two *miro* genes, as a similar distribution was observed in the triple knockouts (Fig. 1). The similar phenotypes between the *miro-1* single mutant and the *miro-1/2/3* triple mutant indicate that *miro-1* is the major functional ortholog. Further, the different phenotypes of *miro* and *trak* mutants indicate that Miro and TRAK do not have equivalent functions in mitochondria transport, as recent studies have suggested (Lopez-Domenech et al., 2018; Zhao et al., 2021). Overall, these results indicate that anterograde, kinesin-1-mediated mitochondria traffic can occur in the absence of Miro and TRAK, suggesting the existence of Miro- and TRAK-independent mechanisms that are dependent on kinesin-1.

**Figure 1.**
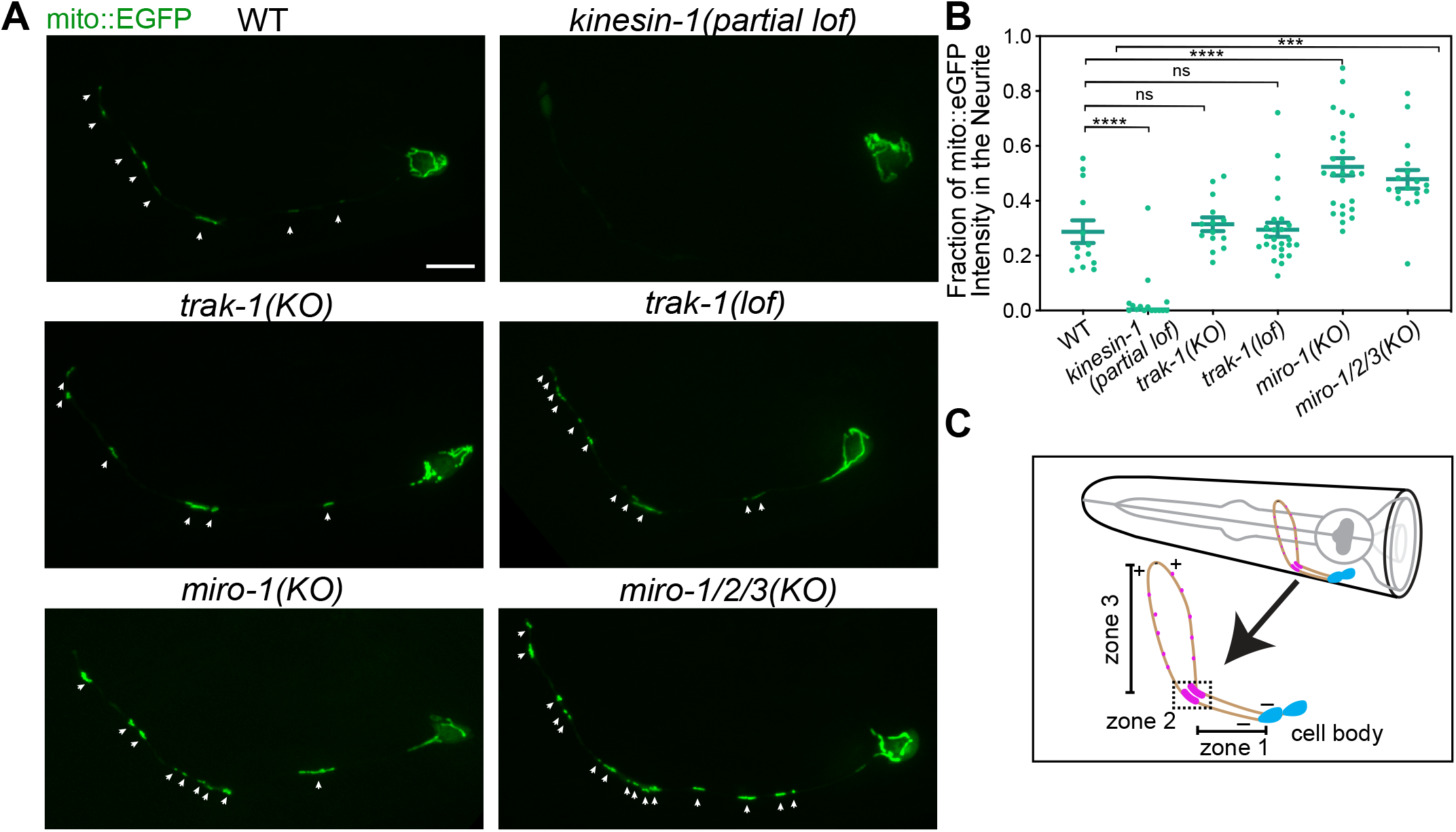
Mitochondria anterograde trafficking requires *kinesin-1*/*unc-116* and does not depend on *miro* or *trak-1*. **(A)** Representative images of mitochondria in AIY in WT and different mutant animals. Arrows indicate mitochondria in the axon. Scale bar = 5 μm. **(B)** Quantification of mitochondria in the axon in WT and different mutant animals. Error bars represent SEM. *** p < 0.001, **** p < 0.0001, ns = not significant (One-way ANOVA with multiple comparisons). **(C)** Schematic diagram of the AIY neurons in the *C. elegans* head. The neurite is composed of 3 zones. Magenta dots in zone 2 and zone 3 indicate clustering of presynaptic vesicles. Microtubule polarity is indicated by + and -.

### Axonal Retrograde Transport is Blocked in *miro-1* Mutants

To understand how mitochondria accumulate in neurites that lack Miro, we performed *in vivo* time-lapse imaging and examined whether anterograde and/or retrograde mitochondria trafficking are affected in *miro-1/2/3* mutants. We imaged larval stage 4 (L4) animals, an age at which AIY neurons have matured and are fully functional. We noticed infrequent mitochondria trafficking events, consistent with the findings that mitochondria motility is less dynamic *in vivo* and in mature neurons (Lewis et al., 2016; Smit-Rigter et al., 2016) (Movie S1). To minimize phototoxicity, we took images every 1 min (as we did not observe significantly more trafficking events with shorter time intervals). To our surprise, mitochondria transport in both directions was dramatically reduced in *miro-1/2/3* mutants. In control animals, we observed an average of 3 mitochondria trafficking events per hour. By contrast, we observed an average of fewer than 1 trafficking event per hour in *miro-1/2/3* mutants, with significantly shorter displacement compared with the control animals in both directions (Fig. S2, Movie S1 and S2). Thus, mitochondria are nearly immobile in *miro* mutants during the window of timelapse-imaging (about 1 hour), precluding analysis of detailed trafficking parameters. These results suggest that axonal mitochondria in *miro* mutants may ‘creep’ into axons at rates too slow to detect by live imaging; alternatively, axonal mitochondria in *miro* mutants might arise from local biogenesis.

To determine if the mitochondria found in AIY neurites in *miro* mutants arise from local biogenesis, we examined overall mitochondria mass during development. We focused on the *miro-1* single mutant since it showed similar axonal mitochondria distribution as *miro-1/2/3* mutants. We used total fluorescence of our mitochondria marker to estimate relative mitochondria mass and normalized this measurement to that of the control larval stage 1 (L1) animals. Using this assay, we found that there is no significant difference in mitochondria mass between control and *miro-1* mutants at any developmental stage. In both control and *miro-1* mutants, total mitochondria mass increases ∼ 40% from the L1 to the larval stage 2 (L2) but does not change significantly from L2 to L4 (Fig. 2A). Next, we examined mitochondria mass specifically in neurites. In control animals, normalized neurite mass increased dramatically from L1 to L2 (0.12 vs. 0.25, P=0.009), and did not change significantly from L2 to L4 (0.25 vs 0.31, P=0.79). Strikingly, *miro-1* mutants had similar neurite mitochondria as L1s but accumulated slightly more neurite mitochondria as L2s (L1: 0.1, L2: 0.37, P<0.0001) and dramatically more neurite mitochondria as L4 (0.77, P<0.0001 compared with L2) (Fig. 2B). The increase in neurite mitochondria from L2 to L4 in *miro-1* mutants was accompanied by a corresponding loss of mitochondria in the cell body (Fig. 2C), consistent with the overall stability of mitochondria total mass (Fig. 2A). Overall, these results suggest that there is an increased net flux of mitochondria in *miro-1* mutants from the cell body into the AIY neurite—in the anterograde direction.

**Figure 2.**
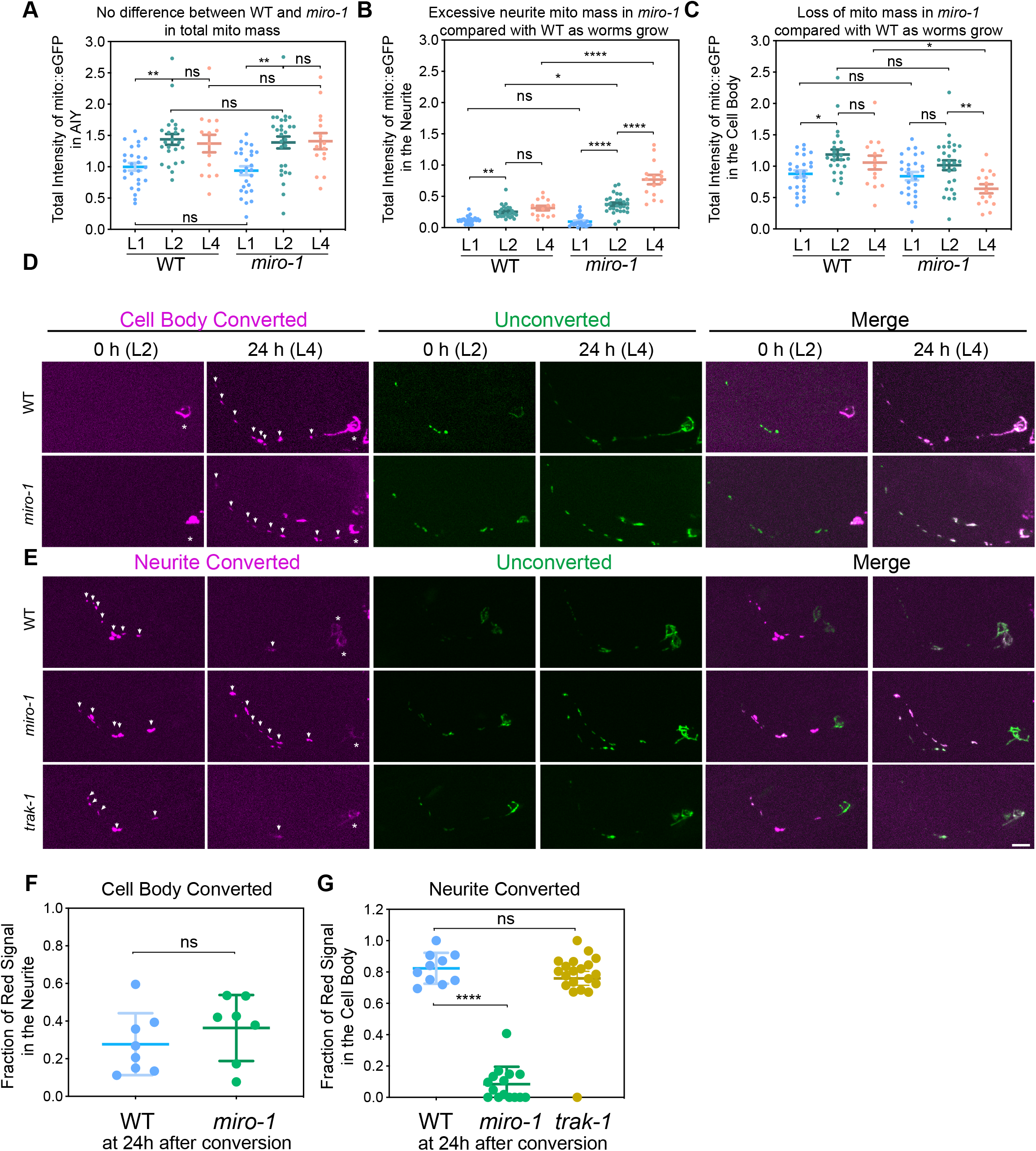
Mitochondria retrograde trafficking requires *miro-1*. **(A-C)** Quantification of mitochondria mass (indicated by the total intensity of mito:EGFP) in WT and *miro-1* mutant animals at larval stage 1, 2 and 4 stages (L1, L2 and L4, respectively). Intensity is normalized to that of the whole AIY neuron in WT L1 animals. Error bars represent SEM. * p < 0.05, ** p < 0.01, **** p < 0.0001 (One-way ANOVA with multiple comparisons). **(D-E)** Representative images of the mito::dendra2 photo-conversion assay. Arrows and asterisks indicate mitochondria in the neurite and in the cell body with converted dendra2 signal, respectively. Scale bar = 5 μm. **(F-G)** Quantification of the mito::dendra2 photo-conversion assay. Left panel: red fluorescence intensity in the neurite/total red fluorescence intensity at 24 hours after the cell body is photoconverted. Right panel: red fluorescence intensity in the cell body/total red fluorescence intensity at 24 hours after the neurite is photoconverted. Error bars represent SEM. **** p < 0.0001, ns = not significant (One-way ANOVA with multiple comparisons).

To confirm and extend this result, we used a photoswitchable fluorophore to perform a ‘pulse-chase’ experiment on mitochondria traffic, enabling mitochondria flux to be assessed at much longer time scales than is possible by continuous live imaging. This assay is powerful in that it integrates rare or short trafficking events over a long period of time. We expressed mitochondria matrix-targeted photoconvertible fluorescent protein Dendra2, and photoconverted mitochondria either in the cell body or in the neurite at L2 stage and imaged 24 hours later (at L4 stage). Mitochondria photoconverted in the cell body showed anterograde flux to the neurite both in control and *miro-1* mutant animals, confirming that MIRO-1 independent anterograde flux moves mitochondria into the neurite (Fig. 2D & F). By contrast, mitochondria photoconverted in the neurite were retained after 24 hours in *miro-1* mutants, while 80% were trafficked back to the cell body in control animals (Fig. 2E & G). These results indicate that MIRO-1 is essential for mitochondria retrograde movement, both on short timescales (Fig. S2) and for long-term flux (Fig. 2E & G). Interestingly, although in some cases Milton/TRAK acts as the adaptor between Miro and dynein (van Spronsen et al., 2013), we found that mitochondria photoconverted in the neurite in *trak-1* mutants were transported back to the cell body similar to controls. This suggests that TRAK-1 is not required for long-term retrograde flux in AIY, unlike Miro (Fig. 2E & G). Overall, our photoconversion experiments are consistent with our mitochondria mass experiments, and indicate that in *miro* mutants mitochondria traffic is anterogradely biased in AIY, resulting in axonal accumulation and cell body depletion over time.

Together, our results clarify the function of Miro *in vivo* in *C. elegans.* Miro is important for robust mitochondria movements, both anterograde and retrograde. However, in the absence of Miro, mitochondria flux persists in the anterograde direction, while retrograde flux is completely blocked.

### Endogenous RIC-7 Localizes at the Leading End of Mitochondria that Undergo Anterograde Trafficking in the Axon

Unlike *miro* mutants, but similar to *kinesin-1* mutants (Fig. 1), *ric-7* mutants lack mitochondria in neuronal processes (Rawson et al., 2014). However, it was not previously determined whether this phenotype is specific to the axon, or if it occurs in both the axon and the dendrite. We found that mitochondria are stuck in the cell body in AIY neurons in *ric-7* mutants (Fig. 3A & D), but these neurons are unipolar and lack classic dendrites. As microtubule polarity is different in axons vs. dendrites, we sought to test if *ric-7* is required for general mitochondria localization, or if it is specific for transport into axons or dendrites. Therefore, we investigated the role of *ric-7* in mitochondria transport in the DA9 neuron. The DA9 neuron is a bipolar excitatory motor neuron, with a dendrite in the ventral nerve cord and an axon that crosses the body and joins the dorsal nerve cord, where it synapses on the body wall muscle (Fig 3C). In DA9, microtubules are plus-end-out in the axon and predominantly minus-end-out in the dendrite (Yan et al., 2013). In DA9, axonal (plus-end directed) mitochondria traffic has been shown to depend entirely on kinesin-1, but only partially on MIRO-1 (Zhao et al., 2021).

**Figure 3.**
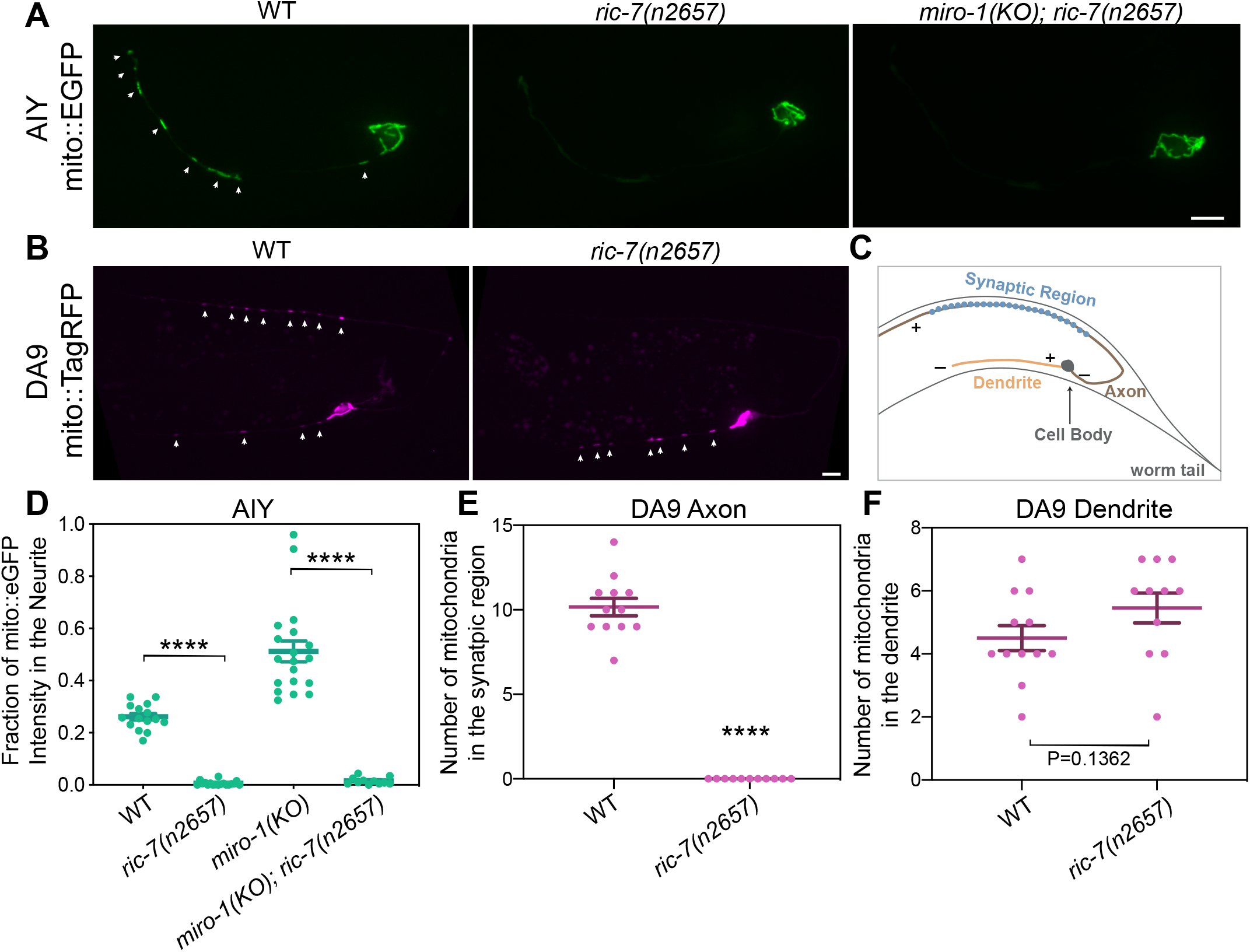
RIC-7 is required for axonal mitochondria localization. **(A)** Representative images of mitochondria in AIY in WT, *ric-7* and *miro-1; ric-7* mutant animals. Arrows indicate mitochondria. Scale bar = 5 μm. **(B)** Representative images of mitochondria in DA9 in WT and *ric-7* mutant animals. Arrows indicate mitochondria. Scale bar = 5 μm. **(C)** Diagram of DA9 neuron in the worm tail. Only the region shown on the micrographs in (B) is depicted. Microtubule polarity is indicated by + and -. **(D-F)** Quantification of mitochondria in AIY neurite, DA9 axon and DA9 dendrite. Error bars represent SEM. **** p < 0.0001 (One-way ANOVA with multiple comparisons for D, t-test for E and F).

We found that mitochondria are absent from the DA9 axon in *ric-7* mutants but are still present (and actually slightly increased) in the dendrite, indicating that *ric-7* is specifically required to transport mitochondria into axons (Fig. 3B, E & F). We previously observed increased net anterograde flux of mitochondria in *miro* mutants (Fig. 2B). Therefore, we tested whether loss of Miro might bypass the requirement for *ric-7* in anterograde flux. We found that regardless of the presence or absence of Miro, mitochondria are absent from the AIY neurite in *ric-7* mutants (Fig. 3D). Thus, long-term anterograde flux in AIY in the absence of MIRO-1 is also RIC-7 dependent. The AIY neurite and the DA9 axon are similar in that both have plus end-out microtubules, while the DA9 dendrite has predominantly minus end-out microtubules. Together, these results indicate that RIC-7 is specifically essential for plus end-directed movement of mitochondria—that is, movement that is mediated by the kinesin-1 motor.

To better understand how RIC-7 regulates mitochondria trafficking, we used the Native and Tissue-Specific Fluorescence (NATF) approach (He et al., 2019) to visualize the endogenous localization of RIC-7 in the DA9 neuron. Briefly, CRISPR was used to insert 7xGFP11 at the endogenous C-terminus of RIC-7, and the complementary GFP1-10 fragment was overexpressed in DA9. This tagging approach does not affect RIC-7’s function as mitochondria localization is not altered (Fig. S3A). Hereafter, we refer the endogenous RIC-7 molecules labeled by reconstituted split-GFP as RIC-7::7xSplitGFP. We found that RIC-7::7xSplitGFP formed puncta that are colocalized with mitochondria in the dendrite and in the axon (Fig. 4A, B). Nearly all RIC-7 puncta colocalized with mitochondria, although we cannot exclude that some RIC-7 is independent of mitochondria below our detection threshold. By contrast, not every mitochondrion was associated with a visible RIC-7::7xSplitGFP puncta. We found that the percentage of mitochondria associated with RIC-7::7xSplitGFP is significantly higher in the axon than in the dendrite (80% in axons; 45% in dendrites; p < 0.0001) (Fig. 4E). Thus, like overexpressed RIC-7 (Rawson et al., 2014), endogenous RIC-7 is highly localized to mitochondria, particularly in axons.

**Figure 4.**
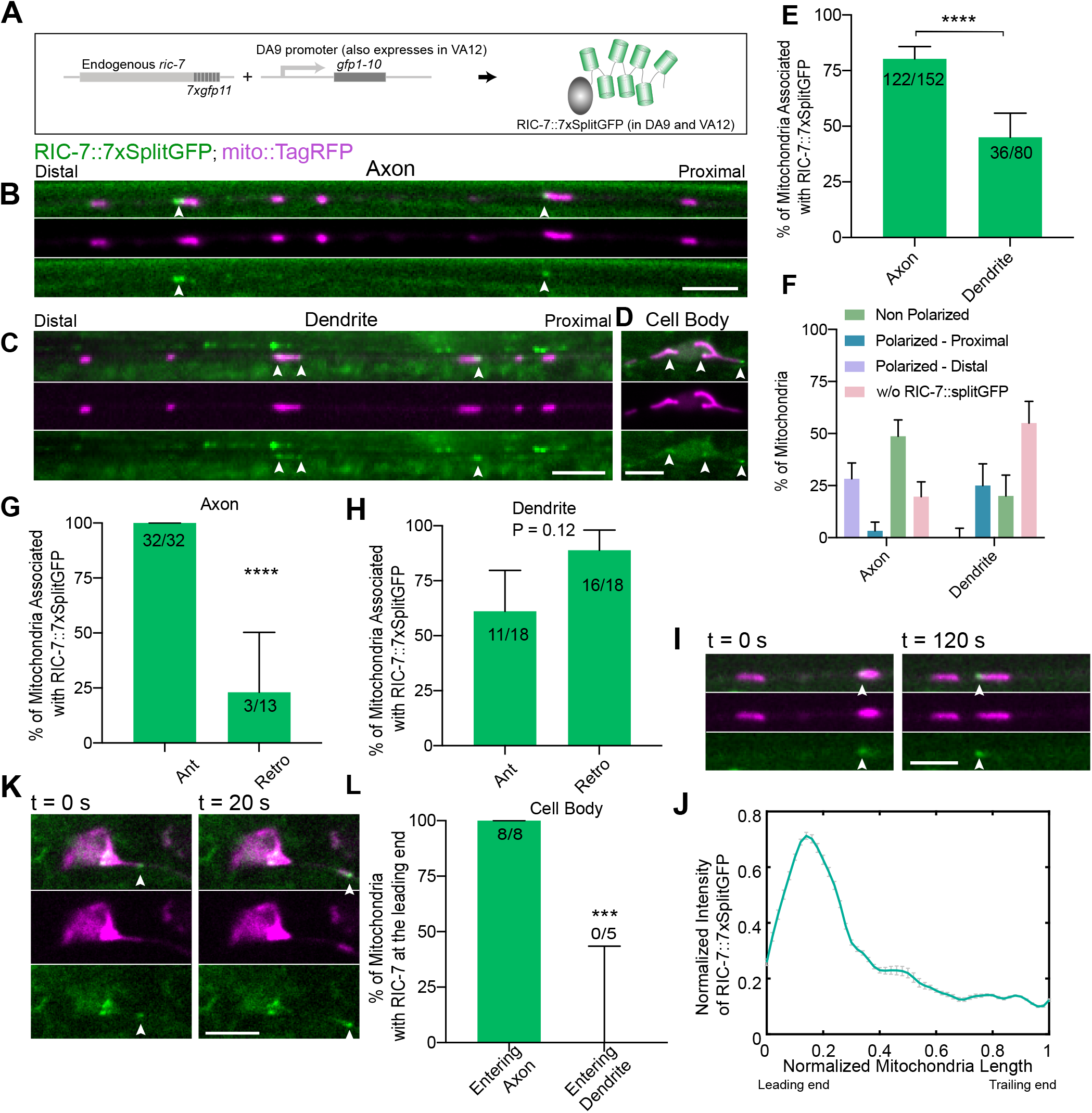
Endogenous Localization of RIC-7. **(A)** Labeling scheme of endogenous RIC-7. **(B-D)** Representative images showing RIC-7::7xSplitGFP puncta associated with mitochondria both in the processes and in the cell body. Note that RIC-7 is also labeled with 7xSplitGFP in VA12, the axon of which is visible next to the dendrite of DA9. Arrowheads indicate RIC-7::7xsplitGFP puncta in DA9. Scale bars = 5 μm. **(E)** Percentage of mitochondria associated with RIC-7::7xsplitGFP puncta in the axon and in the dendrite, quantified from still images. Number of mitochondria counted are indicated at the top of each bar. Error bars represent 95% confidence intervals. **** p < 0.0001 (Fisher’s exact test). **(F)** Percentage of mitochondria associated with RIC-7::7xsplitGFP based on its polarization status from the same dataset as shown in (E). Error bars represent 95% confidence intervals. **(G)** Percentage of moving mitochondria associated with RIC-7::7xsplitGFP puncta in the axon. Number of mitochondria counted are indicated at the top of each bar. Error bars represent 95% confidence intervals. **** p < 0.0001 (Fisher’s exact test). **(H)** Percentage of moving mitochondria associated with RIC-7::7xsplitGFP puncta in the dendrite. Number of mitochondria counted are indicated at the top of each bar. Error bars represent 95% confidence intervals. p value = 0.12 (Fisher’s exact test). **(I)** Representative images of anterogradely trafficking mitochondria in the axon at two different timepoints, with RIC-7::7xSplitGFP accumulating at the leading end, which is indicated by the arrowhead. Scale bar = 2 μm. See Movie S4. **(J)** Normalized intensity of RIC-7::7xsplitGFP averaged across different timepoints along the length of mitochondrion shown in Movie S4. Leading end is on the left and trailing end on the right. Error bars represent SEM. **(K)** Representative images of mitochondria entering the axon, with RIC-7::7xsplitGFP accumulating at the leading end, which is indicated by the arrowhead. Scale bar = 5 μm. See Movie S5. **(L)** Percentage of mitochondria exiting the cell body with RIC-7::7xsplitGFP accumulation at the tip. Number of mitochondria counted are indicated at the top of each bar. Error bars represent 95% confidence intervals. *** p < 0.001 (Fisher’s exact test).

Since RIC-7 is required for mitochondria localization specifically in axons (Fig. 3), we speculated that it might associate with the pool of mitochondria that are mobile in the anterograde direction in the axon. To test this idea, we performed time-lapse imaging and examined the association of RIC-7 with mobile mitochondria in axons and dendrites. As in AIY, mitochondria trafficking events are rare in DA9 - only ∼ 13% of mitochondria are mobile in the axon during the window of imaging (10 min per movie, 10% for anterograde, 3.3% for retrograde, Fig. S3B). We observed that mitochondria moving anterogradely in the axon (plus end-directed) invariably (100%, 32/32) associate with RIC-7::7xSplitGFP puncta. Retrograde mitochondria in the axon typically have no apparent RIC-7, although some do (23%, 3/13) (Fig. 4G, Movie S4). By contrast, in the dendrite, this localization bias is eliminated (Fig. 4H). The strict requirement for RIC-7 in mitochondria axonal localization, and the strong association of RIC-7 with mitochondria undergoing axonal anterograde movement, together suggest that RIC-7 acts on mitochondria to enable their anterograde traffic in the axon.

Interestingly, we observed that the precise localization of RIC-7 on individual mitochondria is sometimes polarized - RIC-7 appears highly localized to one end (Fig. 4A, B). We found that 28% of all axonal mitochondria have RIC-7 puncta at their distal end, closer to the axon tip. By contrast, only 3% of all axonal mitochondria have RIC-7 puncta at their proximal end, closer to the cell body. This polarized localization pattern is reversed in the dendrite: 0% of all dendritic mitochondria have RIC-7 puncta at their distal end, closer to the dendrite tip, and 25% have RIC-7 puncta at their proximal end (Fig. 4F). Thus, RIC-7 tends to localize to the end of mitochondria that correlates with the direction of the plus end of local microtubules—that is, the leading end for kinesin-1 mediated transport.

We speculated that mitochondria anterograde movement might be correlated with the polarized localization of RIC-7 at the leading edge. We tested this idea using time-lapse imaging. We observed that anterograde trafficking mitochondria in the axon often have RIC-7::7xSplitGFP puncta at their leading end (Fig. 4I & J, Movie S3 and S4). However, although we could tell that all anterogradely-moving mitochondria are associated with RIC-7 (Fig. 4G), some mitochondria are too small to allow us to resolve the distribution of RIC-7 on mitochondria. Therefore, to better quantify the relationship between RIC-7 leading end localization and mitochondria trafficking, we examined mitochondria exiting the cell body. As previously shown, mitochondria exiting the cell body usually first extend into the axon, followed by fission and trafficking (Zhao et al., 2021) (Fig. 4G). Strikingly, as mitochondria extend into the axon, we found that RIC-7 accumulates at the mitochondria tip, leading its way into the axon (Fig. 4K & L, Movie S5). This happened to all mitochondria we observed entering the axon (8/8, Fig. 4L). Importantly, we did not observe RIC-7 accumulation at the leading end of mitochondria entering the dendrite (Fig. 4L and Movie S6). Overall, these data suggest that RIC-7 acts at the leading end of mitochondria to allow kinesin-1-mediated movement.

### Endogenous Localization of Kinesin-1

To investigate the cell biology of kinesin-1 relative to RIC-7 and mitochondria, we labeled endogenous kinesin-1/UNC-116 using the NATF approach (He et al., 2019) and visualized its distribution along with mitochondria in DA9 neurons (Fig. 5 A & B). Although kinesin-1 is known to transport mitochondria, how it is distributed on motile mitochondria has not been reported previously. We found that insertion of 3xGFP11 at the C-terminus doesn’t interfere with the function of kinesin-1, as these animals show normal body locomotion, mitochondria localization and trafficking (Fig. 5A, Movie S7). (We note that 7xGFP11 did interfere with kinesin-1/UNC-116 function; we hypothesize that the larger 7x tag may interfere with important interactions.) Endogenous kinesin-1::3xsplitGFP is diffuse in the cell body and throughout the axon and dendrite, similar to the localization of the endogenous kinesin-1 in Drosophila (Kelliher et al., 2018). This diffuse distribution is also consistent with the idea that most kinesin-1 molecules are in the autoinhibited conformation (Blasius et al., 2007). In addition to this diffuse pool, kinesin-1::3xsplitGFP also forms visible puncta in the cell body and in the axon (Fig. 5B).

**Figure 5.**
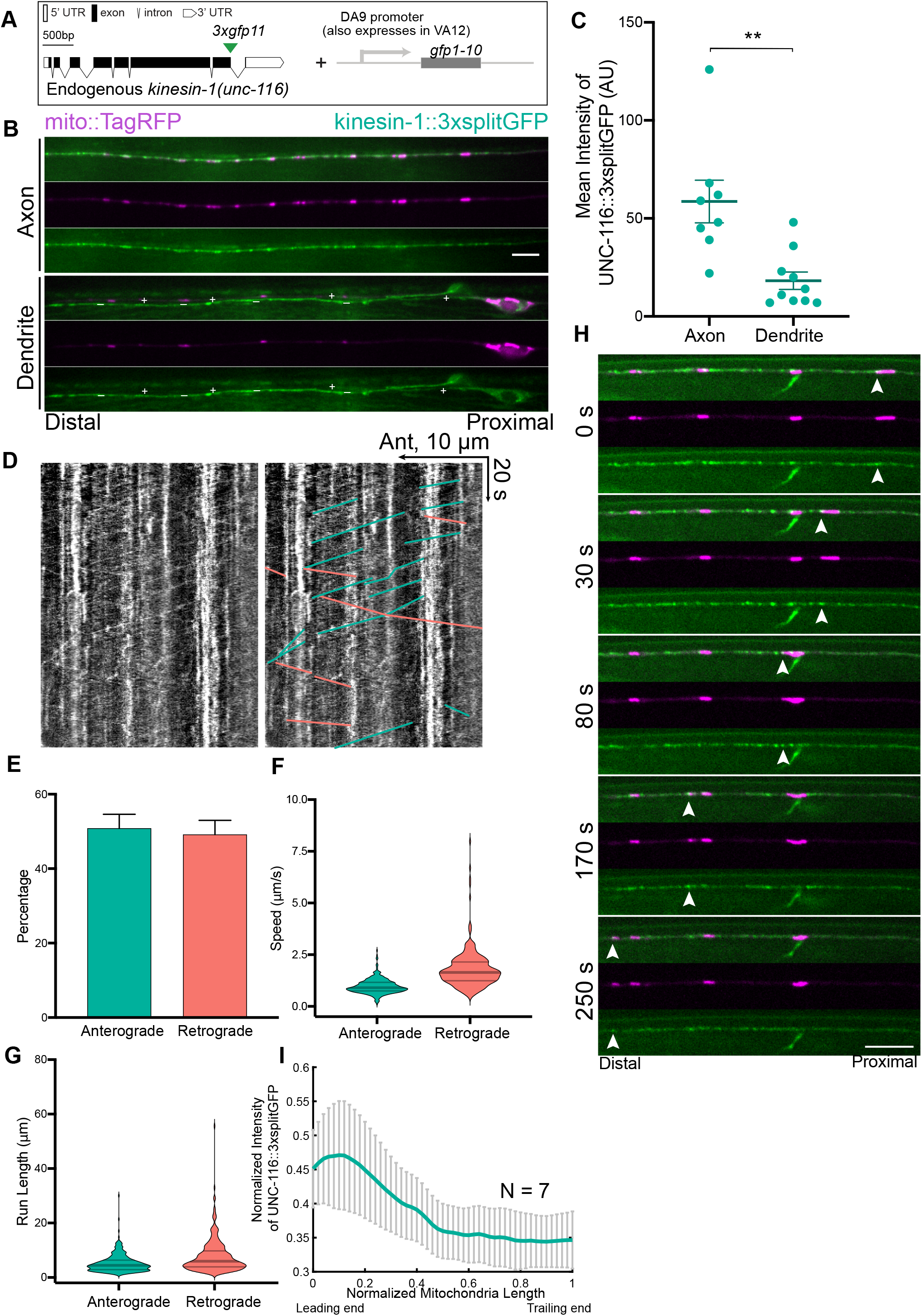
Endogenous Localization of kinesin-1/UNC-116. **(A)** Labeling scheme of endogenous kinesin-1/UNC-116. **(B)** Representative images showing the localization of kinesin-1::3xSplitGFP (UNC-116::3xSplitGFP) in the axon and in the dendrite. + indicates the DA9 dendrite, whereas - indicates the VA12 axon. Note that the dendritic kinesin-1::3xsplitGFP is much less abundant. Scale bars = 5 μm. **(C)** Quantification of kinesin-1::3xSplitGFP intensity in DA9 axon and dendrite. ** p < 0.01 (t test). **(D)** Kymograph showing anterograde and retrograde trafficking events of kinesin-1::3xSplitGFP in the axon with (left) or without (right) manual traces. Green: anterograde; salmon: retrograde. **(E-G)** Quantification of anterograde and retrograde trafficking events of kinesin-1::3xSplitGFP in the axon. (E) Percentage of anterograde and retrograde trafficking events, respectively. Error bars represent 95% confidence intervals. (F-G) Speeds and run lengths of anterograde and retrograde trafficking events, respectively. Violin plots with median and quartiles are shown. **(H)** Representative images showing that kinesin-1::3xSplitGFP is enriched at the leading end of a mitochondrion moving anterogradely. Arrow heads highlight the accumulation of kinesin-1::3xSplitGFP. **(I)** Normalized intensity of kinesin-1::3xsplitGFP along mitochondria length averaged across different timepoints during anterograde trafficking, and then averaged across 7 different mitochondria. Leading end is on the left and trailing end on the right. Error bars represent SEM.

Overall, kinesin-1::3xsplitGFP is more abundant in the axon than in the dendrite, consistent with its plus end-directed motor activity (Fig. 5B & C). Live imaging shows that kinesin-1::3xsplitGFP undergoes extensive anterograde and retrograde trafficking in the axon, with no significant directional bias (Fig. 5D & E). The average speed of anterograde trafficking is 0.97 μm/s, consistent with the reported kinesin-1 velocity measured *in vitro* (Schnitzer et al., 2000; Verbrugge et al., 2009)(Fig. 5F). The average speed of retrograde trafficking is 1.78 μm/s. These retrograde events presumably reflect the binding of kinesin-1 to cargoes that are transported back to the cell body by dynein, since previous studies have demonstrated that kinesins and dynein can bind to the same cargo in axons (Encalada et al., 2011; Hendricks et al., 2010; Szpankowski et al., 2012). The observed average run lengths of anterograde trafficking and retrograde trafficking are 5.2 μm and 7.8 μm, respectively (Fig. 5G).

Next, we examined the colocalization of kinesin-1::3xsplitGFP with moving mitochondria. Unlike RIC-7, a substantial fraction of axonal and dendritic kinesin-1/UNC-116 is not associated with mitochondria. Nevertheless, 80% of mitochondria (8/10) that are trafficked anterogradely in the axon have visible kinesin-1::3xsplitGFP puncta associated with them. The remaining mitochondria were surrounded by diffuse kinesin-1::3xsplitGFP. Similar to the localization of RIC-7, kinesin-1::3xsplitGFP is enriched at the leading edge of anterogradely moving mitochondria, albeit to a lesser extent (Fig. 5H & I, Movie S7). We suspect that the abundant kinesin-1 that is not associated with mitochondria reduces our ability to assess its fine localization on mitochondria, possibly accounting for the lesser degree of polarized localization. Overall, these experiments provide the first view of kinesin-1’s localization on motile mitochondria *in vivo* and demonstrate that kinesin-1/UNC-116 has polarized localization to the leading edge of anterogradely trafficking mitochondria, similar to RIC-7.

Given that kinesin-1 and RIC-7 are both localized to the leading end of anterogradely trafficking mitochondria, we sought to examine whether the polarized localization of RIC-7 is dependent on kinesin-1/UNC-116. Mitochondria transport into the axon is blocked in *kinesin-1* mutants, making it impossible to examine RIC-7 localization on axonal mitochondria in this background. Therefore, we performed acute cell-specific knockdown of kinesin-1 using the auxin degron system (Fig. 6A). After 5 hours of auxin treatment, mitochondria in the dendrite accumulate at the distal tip, indicating that kinesin-1 is degraded, allowing dynein-mediated transport to predominate (Fig. 6B). By contrast, the number of mitochondria in the synaptic region in the axon stayed constant (Fig. 6B & D). This suggests that *in vivo*, synaptic mitochondria are anchored by certain mechanisms that allow them to resist transport imbalances, as previously suggested (Gutnick et al., 2019). We used these synaptic mitochondria to assess the localization of RIC-7 in neurons with reduced kinesin-1. We found that the polarized localization of RIC-7 is nearly abolished after kinesin-1 knockdown (control: 27.3%; auxin: 4.5%; p < 0.0001) (Fig. 6B, C & E). We argue that the loss of polarized localization of RIC-7 is not simply a secondary effect caused by the absence of mitochondria anterograde trafficking, as the percentage of mobile mitochondria in normal conditions (10% for anterograde, Fig. S3B) is far less than the percentage of mitochondria with polarized RIC-7 (27-28%, Fig. 4F & Fig. 6E). Rather, this result indicates that kinesin-1 is required for polarized RIC-7 localization on mitochondria. Further, kinesin-1 knockdown increases the percentage of mitochondria without apparent colocalization of RIC-7 (control: 35.2%; auxin: 57.1%; p = 0.0006) (Fig. 6E). The decrease in overall association of mitochondria with RIC-7 might be due to a difference in detection— unpolarized RIC-7 might simply be more difficult to detect. Alternatively, kinesin-1 may help stabilize the localization of RIC-7 on mitochondria. Taken together, these data indicate that kinesin-1 is required for polarized RIC-7 localization on mitochondria, and that RIC-7 binding to mitochondria is largely independent of kinesin-1.

**Figure 6.**
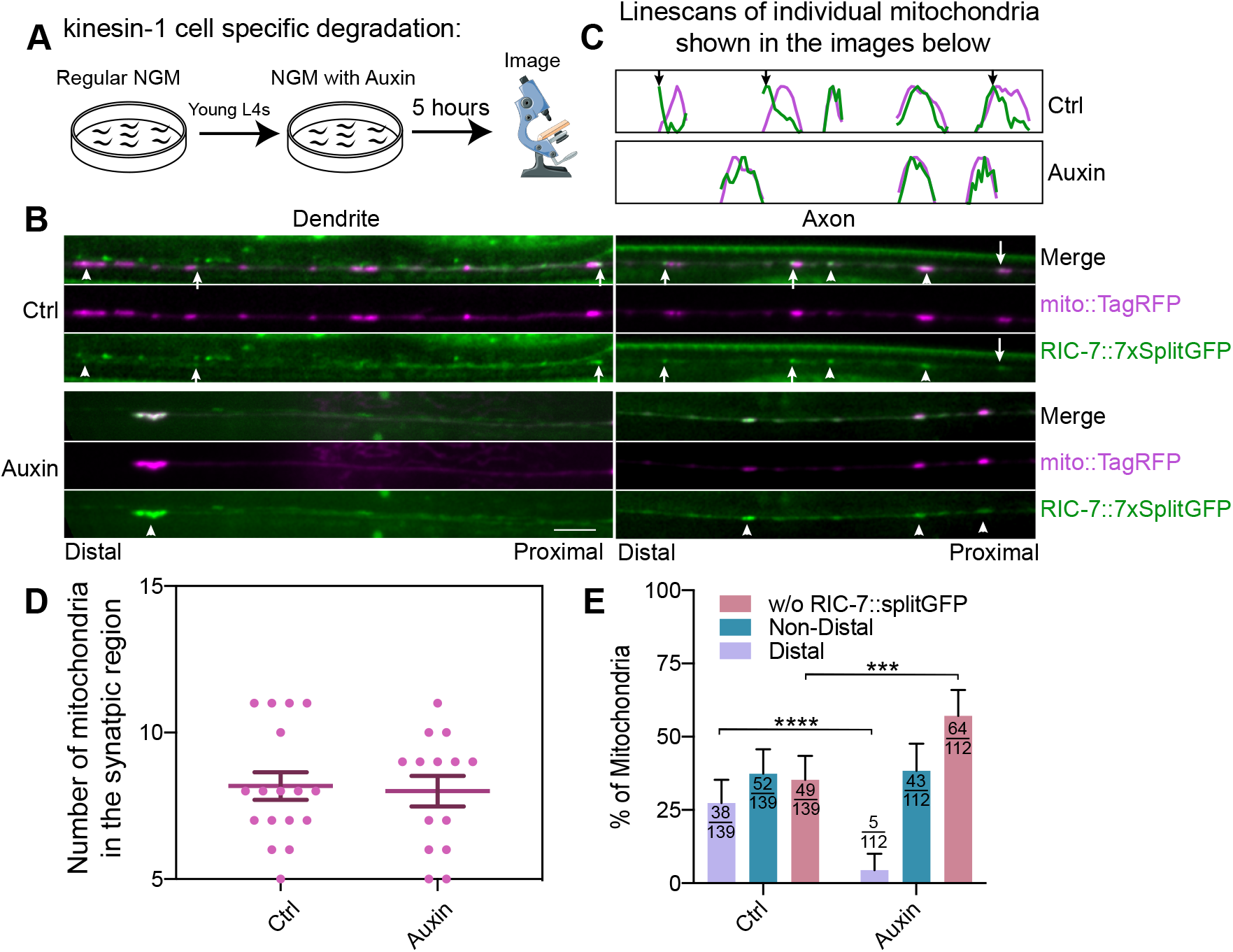
Kinesin-1 controls the polarized localization of RIC-7. (A) Schematic representation of cell specific knockdown of kinesin-1. (B) Representative images of control and *kinesin-1*/*unc-116* knockdown conditions in the dendrite (left) and in the axon (right). Note the accumulation of mitochondria at the distal tip of the dendrite under auxin treatment, indicating kinesin-1 has been efficiently degraded. Arrow heads point to unpolarized RIC-7::7xsplitGFP puncta, while arrows indicate the polarized ones. (C) Linescans of individual mitochondria in the axons shown in (B). Green traces represent the intensity of RIC-7::7xsplitGFP whereas magenta traces represent that of mito::TagRFP. Arrows indicate the polarized localization of RIC-7::7xSplitGFP. (D) The number of axonal mitochondria does not change after 5 hours of auxin treatment. Error bars represent SEM. (E) Quantification of the percentage of mitochondria based on RIC-7::7xsplitGFP’s polarization status. Error bars represent 95% confidence intervals. *** p < 0.001, **** p < 0.0001 (Fisher’s exact test)

In summary, we find both RIC-7 and kinesin-1 accumulate at the leading end of anterogradely trafficking mitochondria in the axon, and kinesin-1/UNC-116 controls the localization of RIC-7, supporting the hypothesis that kinesin-1 and RIC-7 work together to drive mitochondria movement toward the microtubule plus end.

### MIRO-1 and MTX-2 Control the Localization of RIC-7

Along with MIRO-1, MTX-2 has been identified as an important regulator of mitochondria trafficking. It is proposed that MTX-2 and MIRO-1 form the adaptor core for both kinesin-1 and dynein motors, with MTX-2 playing the major role (Zhao et al., 2021). Similar to the findings in DA9 and PVD neurons shown in Zhao et al., 2021, the AIY neuron has a reduced number of mitochondria in *mtx-2* mutants, and a complete loss of mitochondria in *mtx-2; miro-1* double mutants (Fig. 7A & B). We find that RIC-7 is required for mitochondria transport into axons, but not into dendrites (Fig. 3), in contrast to the Miro-1/metaxin2 combination, which is required for transport into both axons and dendrites (Zhao et al., 2021). Thus, the potential relationship between RIC-7 and Miro/Metaxin is unclear.

**Figure 7.**
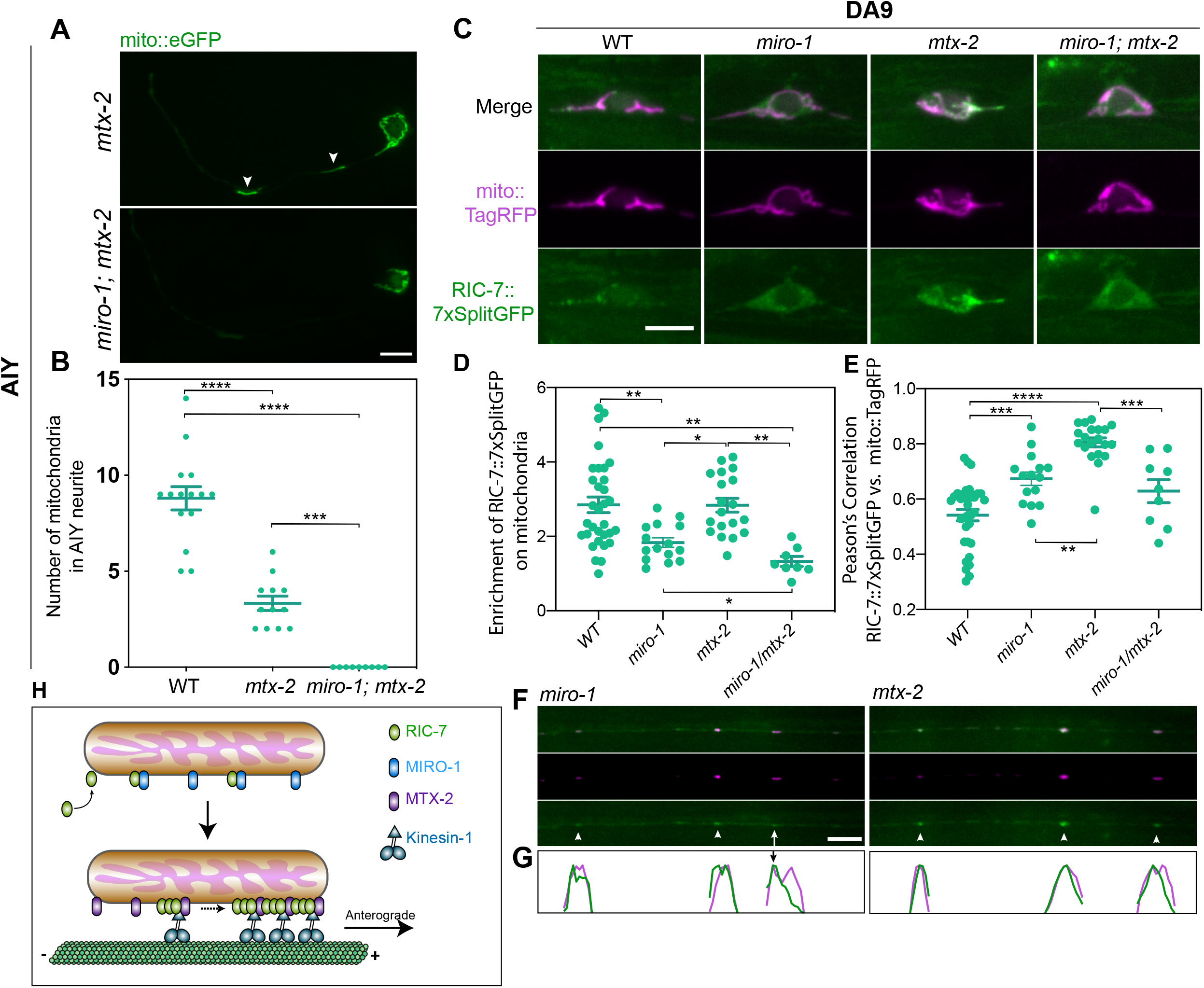
MIRO-1 and MTX-2 regulate the endogenous localization of RIC-7. **(A)** Mitochondria distribution in AIY neurons in *mtx-2* mutant and *mtx-2; miro-1* double mutants. Arrow heads indicate mitochondria. Scale bar = 5 μm. **(B)** Quantification of mitochondria in the AIY neurite in WT, *mtx-2* mutant and *mtx-2; miro-1* double mutants. Error bars represent SEM. *** p < 0.001, **** p < 0.0001 (One-way ANOVA with multiple comparisons). **(C)** Endogenous RIC-7::7xsplitGFP localization in the DA9 cell body of WT, *miro-1*, *mtx-2* and *miro-1; mtx-2* mutants. Scale bar = 5 μm. **(D)** Quantification of the enrichment of RIC-7::7xSplitGFP on mitochondria in WT, *miro-1*, *mtx-2* and *miro-1; mtx-2* mutants. * p < 0.05, ** p < 0.01 (One-way ANOVA with multiple comparisons). **(E)** Pearson’s correlation between RIC-7::7xSplitGFP and mito::TagRFP in WT, *miro-1*, *mtx-2* and *miro-1; mtx-2* mutants. *** p < 0.001, **** p < 0.0001 (One-way ANOVA with multiple comparisons). **(F)** Endogenous RIC-7::7xsplitGFP localization in the DA9 axon of *miro-1* and *mtx-2* mutants. Arrow heads point to unpolarized RIC-7::7xsplitGFP puncta, while arrows indicate the polarized ones. **(G)** Linescans of individual mitochondria in the axons shown in (F). Green traces represent the intensity of RIC-7::7xsplitGFP whereas magenta traces represent that of mito::TagRFP. Arrow indicates the polarized localization of RIC-7::7xsplitGFP. **(H)** Proposed Model for Axonal Anterograde Transport of Mitochondria in *C. elegans*. RIC-7 is recruited to mitochondria by MIRO-1-dependent and independent mechanisms. MTX-2 helps enrich RIC-7 to the leading end of mitochondria, potentially forming a complex with RIC-7 and kinesin-1, which is essential for mitochondria anterograde trafficking.

To test whether Miro and metaxin2 function independently or together with RIC-7 for kinesin-1 mediated mitochondria transport, we characterized the localization of RIC-7 in *miro-1* and *mtx-2* mutants. In the DA9 cell body of *miro-1* mutants, RIC-7 becomes much more cytosolic and less enriched on mitochondria (Fig. 7C). We quantified this distribution in two ways, focusing on mitochondria in the cell body since these are present in all genotypes. First, we quantified the enrichment of RIC-7 on mitochondria by normalizing the intensity of GFP puncta relative to the cytosolic signals (Fig. 7D, see methods). Second, we calculated the pixel-by-pixel Pearson’s Correlation between RIC-7 and mitochondria: the more evenly RIC-7 is distributed, the higher the correlation will be. Using these approaches, we found that RIC-7 is less enriched on mitochondria in *miro-1* mutants (1.8 fold) than in controls (2.8 fold, Fig. 7D), and is significantly more evenly distributed (less punctate) (Fig. 7E). Axonal mitochondria in *miro-1* mutants still have RIC-7 signals, indicating that there is a MIRO-1-independent mechanism for RIC-7 to localize to mitochondria (Fig. 7F). Further, RIC-7 is still polarized to the distal end of some axonal mitochondria in *miro-1* mutants, albeit to a lesser degree compared with WT (Fig. 7G). These results—together with our functional data (Fig. 2, Movie S2)—indicate that MIRO-1 is important for RIC-7’s normal localization and function, but that some of RIC-7’s localization and function remain even in the absence of MIRO-1.

In the cell body of *mtx-2* mutants, RIC-7::7xsplitGFP is as enriched on mitochondria as in control animals (2.8 fold enrichment). However, unlike in WT where it forms discrete puncta, RIC-7 becomes extremely dispersed over the entire mitochondria, to a greater extent than in *miro-1* mutants (Fig. 7C, E). In the axon, although RIC-7::7xsplitGFP is still localized to mitochondria the polarized localization at the leading end is absent (Fig. 7F & G, Movie S8). Thus, MTX-2 has a primary role in enabling RIC-7 localization to mitochondria subdomains.

We also examined RIC-7 distribution in *miro-1; mtx-2* double mutants. In these mutants, mitochondria are trapped in the cell body and fail to traffic into both axon and dendrite (Zhao et al., 2021). Thus, distribution of RIC-7 on axonal mitochondria cannot be determined. In the cell body, combined effects from single mutants were found: RIC-7 does not form discrete puncta and becomes less enriched (1.3 fold) on mitochondria (Fig. 7C, D, E).

Overall, this analysis suggests that MIRO-1 and MTX-2 have different roles in mediating RIC-7’s mitochondria localization. MIRO-1 is involved in overall localization of RIC-7 to mitochondria, but some RIC-7 can localize to mitochondria even without MIRO-1. By contrast, MTX-2 is required to concentrate RIC-7 at specific loci on mitochondria, functioning together with kinesin-1 to polarize RIC-7 distribution toward the plus-end of microtubules.

### Structure/Function Analysis of RIC-7 Identifies Regions Required for Its Localization and Function

The previous data suggests that RIC-7 has at least two features. First, to localize to mitochondria, partially independently of MIRO-1. Second, to accumulate at the leading end of mitochondria and enable mitochondria anterograde movement. To better understand how RIC-7 functions, we performed a structure/function analysis. The endogenous genomic sequence of the *ric-7* gene spans ∼ 20kb and overlaps with two other genes, which would make structure/function analysis at the endogenous locus difficult. Therefore, we decided to perform this analysis using transgenes expressed as single copy insertions at a standardized genomic locus under the same promoter, so that different transgenes will achieve comparable levels of expression. The *ric-7* gene encodes for 3 different splice variants (Fig. S5B), two of which (*ric-7a* and *ric-7b*) can rescue mitochondria localization defects in *ric-7* mutants (Rawson et al., 2014). We decided to focus on *ric-7b* because this isoform is preferentially expressed compared with *ric-7a* across different neurons (Fig. S5B & C)(Taylor et al., 2021).

The predicted structure of RIC-7B by Alphafold (Jumper et al., 2021; Varadi et al., 2022) shows three distinct domains: a relatively ordered domain at the N-terminus (1-99aa), a largely disordered domain in the middle (100-470aa), and a structured domain at the C-terminus (471-709aa) that is similar to the OTU domain in human otulin, according to HHPred and Dali (Holm et al., 2023; Soding et al., 2005)(Fig. 8A, Fig. S6 and Movie S9). The N-terminal and the middle regions do not resemble any known structures. Based on this information, we expressed deletions of RIC-7, and assessed their localization and abilities to rescue mitochondria localization defects in *ric-7* mutants. To enable visualization of the localization of these truncated proteins, all versions of RIC-7 were tagged at the C terminus with EGFP.

**Figure 8.**
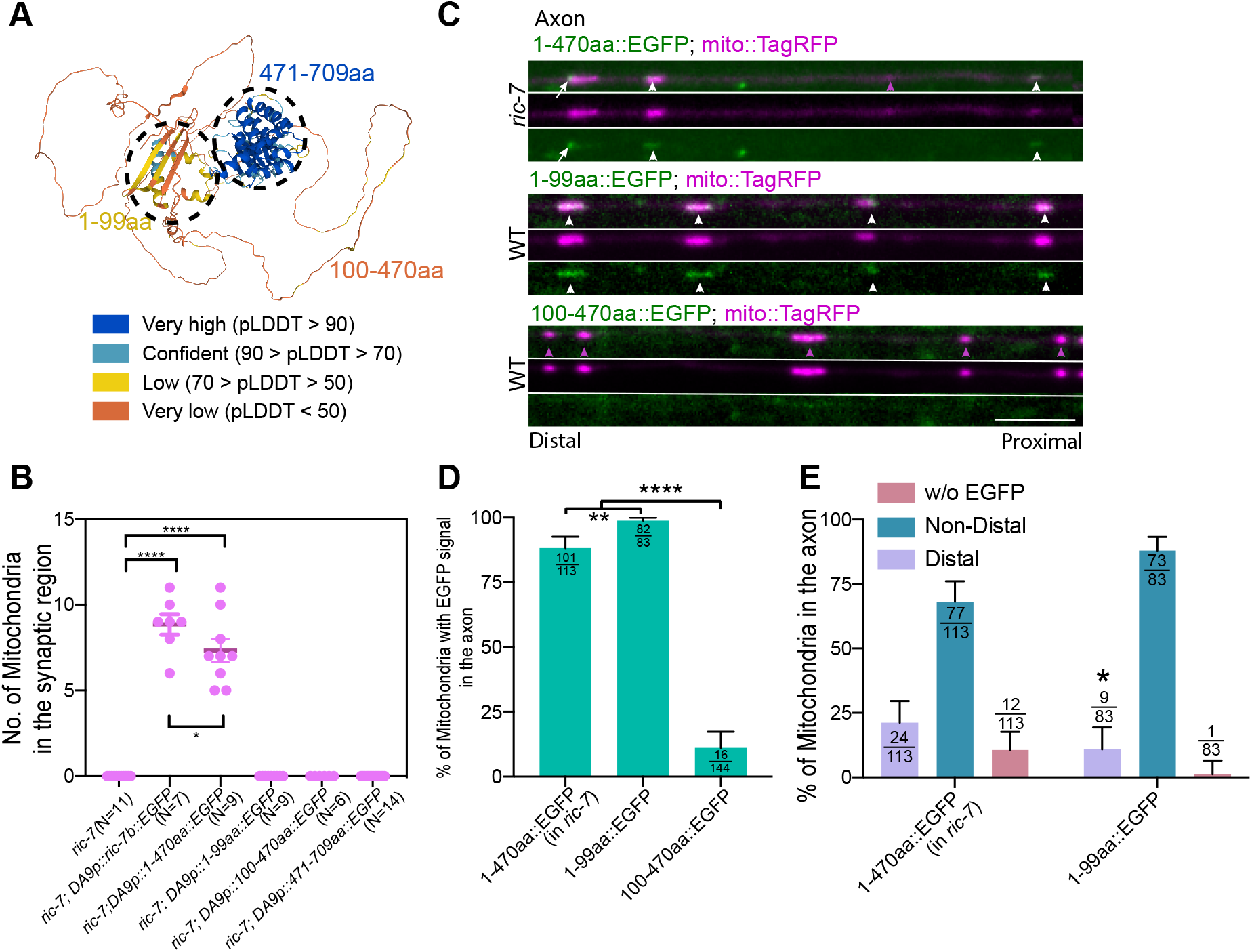
Structure/Function analysis of RIC-7. **(A)** 3D structure of RIC-7b predicted by Alphafold. Model confidence is color coded. The highest confidence region (471-709aa) is structurally similar to human OTU domain. **(B)** Quantification of the ability of different RIC-7 truncations to rescue mitochondria localization defects in *ric-7* mutants. Transgenes were driven by *itr-1* promoter (see methods). *** p < 0.001, **** p < 0.0001 (One-way ANOVA with multiple comparisons). **(C)** Localization of truncated proteins. White arrows highlight the distal end localization of 1-470aa::EGFP; White arrowheads indicate the non-distal localization of truncated proteins; Magenta arrowheads indicate mitochondria that are not associated with the truncated proteins; Scale bar = 5 μm. **(D)** Quantification of the percentage of mitochondria associated with the GFP transgenic fusions. Number of mitochondria counted are indicated at the top of each bar. Error bars represent 95% confidence intervals. ** p < 0.01, **** p < 0.0001 (Fisher’s exact test). **(E)** Quantification of the percentage of mitochondria based on the polarization status of the GFP transgenic fusions. Same dataset as in (D). * p < 0.05 (Fisher’s exact test of Non-distal vs. Distal).

Expression of full-length RIC-7::eGFP restored abundant axonal mitochondria in *ric-7* mutants. Expression of 1-470aa::eGFP (lacking the region similar to OTU domain) also largely rescued mitochondria localization defects in *ric-7* mutants, suggesting that 471-709aa is mostly dispensable for RIC-7’s function. By contrast, expression of the C-terminal OTU domain 471-709aa did not rescue mitochondria localization defects in *ric-7* mutants (Fig. 8B). We also used a stronger pan-neuronal promoter *Prab-3* to drive the expression of either the OTU domain or the human Otulin or OtulinL, none of which could rescue mitochondria distribution defects in *ric-7* mutants (Fig. S7). Consistent with its ability to rescue mitochondria phenotype in *ric-7* mutants, 1-470aa::eGFP forms puncta that are colocalized with mitochondria in the axon, some of which are at the distal end, similar to endogenously-tagged RIC-7 (Fig. 8C, D, E). We further found that expression of either 1-99aa or 100-470aa alone cannot rescue mitochondria localization defects in *ric-7* mutants, indicating that both of these domains are required for RIC-7’s function (Fig. 8B). Interestingly, RIC-7 1-99aa::eGFP colocalizes with mitochondria in both WT and *ric-7* animals (Fig. 8C & D, Fig. S4). In the axon of WT animals, 1-99aa::eGFP is colocalized with 99% of mitochondria; however, it is only enriched to the distal end of 10% of mitochondria, compared with 21% for 1-470aa::eGFP, suggesting that 100-470aa helps RIC-7 form puncta at mitochondria subdomains (Fig. 8E). We surmise that the weak polarized localization of 1-99aa may be dependent on endogenous RIC-7, but we cannot examine 1-99aa localization on axonal mitochondria in the *ric-7* mutant background (as there are no axonal mitochondria). However consistent with this idea, in the *ric-7* mutant background, 1-99aa::eGFP covers the entire mitochondria in the cell body and in the dendrite, unlike the endogenous full length RIC-7, which forms discrete puncta (Fig. S4A & B, Fig. 4C & D). Interestingly, the overall localization of 1-99aa::eGFP to mitochondria is not dependent on MIRO-1, suggesting that 1-99aa contains a mitochondria localization signal that either directly binds to mitochondria or to mitochondria protein(s) other than MIRO-1 (Fig. S4A). In comparison, 100-470aa::eGFP is largely cytoplasmic in WT animals, with only 11% of axonal mitochondria associated with 100-470aa::eGFP signals. In *ric-7* mutant animals, this association is slightly increased (49% of dendritic mitochondria contain eGFP signals), suggesting that 100-470aa can only weakly bind to mitochondria, and is displaced by full-length RIC-7 (Fig. 8C & D, Fig. S4).

In summary, we find that RIC-7 1-470aa is sufficient to support mitochondria transport and that 1-99aa is required for mitochondria localization, while the disordered domain (100-470aa) facilitates its polarized localization on mitochondria.

## Discussion

Mitochondria transport is critical to the maintenance of various neuronal functions. In this study, we used the C. *elegans* AIY neuron and the DA9 neuron as model systems to study mitochondria transport *in vivo*. We characterize RIC-7 as an essential regulator of kinesin-1 based mitochondria trafficking in *C. elegans*. Both kinesin-1 and RIC-7 act at the leading end to drive axonal anterograde mitochondria transport. The N-terminal domain of RIC-7 (1-99aa) targets to mitochondria and the disordered domain facilitates RIC-7’s accumulation at the mitochondria leading end. We find that Miro is important but not essential for mitochondria anterograde transport in the axon, as a Miro-independent and RIC-7 dependent mechanism can move mitochondria anterogradely. RIC-7 partially relies on MIRO-1 to localize to mitochondria and requires MTX-2 to accumulate at the leading end, where RIC-7 promotes kinesin-1-based mitochondria transport (Fig. 7H). We propose that the polarized localization of kinesin-1 and RIC-7 underlies anterograde mitochondria trafficking in the axon.

### RIC-7 functions with Miro and metaxin2 to mediate mitochondria anterograde traffic

Previous studies in multiple model systems showed that loss of function of Miro results in variable phenotypes in mitochondria distribution. While loss of function of Miro leads to dramatic decrease of mitochondria in axons in Drosophila photoreceptors and motor neurons, axonal mitochondria distribution in hippocampal neurons cultured from Miro knockout mice was not altered (Guo et al., 2005; Lopez-Domenech et al., 2016; Stowers et al., 2002). In *C. elegans,* loss of Miro does not cause a change in mitochondria number in touch receptor neurons and PVQ neurons (Ding et al., 2022; Sure et al., 2018). By contrast, in the *C. elegans* DA9 neuron, *miro* loss-of-function leads to reduced mitochondria in the axon (Zhao et al., 2021). This study shows that in the AIY neuron, loss of Miro results in *increased* mitochondria number. What mechanisms might account for these different phenotypes?

We find that in the absence of Miro, axonal mitochondria are largely frozen in place. Further, RIC-7 localization to mitochondria is reduced. One possibility is that the right amount of RIC-7 on mitochondria is essential for processive movements, and without MIRO-1, RIC-7 does not reach this optimal level. In this model, all Miro is doing is to localize RIC-7 to mitochondria. Alternatively, Miro itself may promote kinesin-1-mediated mitochondria transport, such as by strengthening the interaction between kinesin-1 and mitochondria. Nevertheless, over long-time scales, mitochondria can still traffic anterogradely in *miro* mutants, in a process that depends entirely on RIC-7 and kinesin-1. Cell-type-specific differences in this novel form of Miro-independent mitochondria anterograde traffic may account for the variable phenotype observed in Miro mutants. Alternatively, variation in other mechanisms that affect overall mitochondria distribution, such as retrograde traffic or mitophagy, might underlie these differences.

Our data also support the idea that metaxin2 is partially redundant with Miro for anterograde mitochondria traffic. We find that loss of *mtx-2* eliminates mitochondria anterograde traffic in Miro mutants in AIY, consistent with previous results in DA9 and PVD (Zhao et al., 2021). We also find that RIC-7 localization is disrupted in *mtx-2* mutants, failing to form puncta on mitochondria. Nevertheless, some mitochondria still traffic to the axon in *mtx-2* mutants, and this pool is dependent on Miro. Together, these data suggest that RIC-7 is a key point of convergence for anterograde traffic, and that MTX-2 and MIRO-1 work to coordinate RIC-7 function.

We also analyzed Miro’s role in dynein-based trafficking. Using photoconversion, we found that mitochondria retrograde trafficking in axons is completely blocked in *miro-1* mutants. Consistently, Miro loss-of-function also results in mitochondria disappearance from DA9 dendrites and PVD anterior dendrites in *C. elegans* (Zhao et al., 2021). Further, in hippocampal neurons cultured from Miro knockout mice, dendritic mitochondria are dramatically reduced while axonal mitochondria distribution are not altered (Lopez-Domenech et al., 2016). All together, these results highlight Miro’s essential role in dynein-based mitochondria trafficking.

Biochemical and functional analysis suggests that Miro is linked to the dynein/dynactin complex via Milton/ TRAK (Brickley and Stephenson, 2011; van Spronsen et al., 2013) (Zhao et al., 2021). Surprisingly, we found that mitochondria distribution in the AIY neuron is not altered in *trak-1* mutants, and that long-term retrograde trafficking is preserved. These data suggest that either other adaptors may link Miro and the dynein complex in AIY, or that the *C. elegans* Miro can directly bind to the dynein complex. These possibilities await further investigation.

Taken together, our data demonstrate that anterograde mitochondria traffic in *C. elegans* relies on two essential factors, kinesin-1 and RIC-7, which function together with Miro and metaxin.

### Kinesin-1 and RIC-7 Drive Mitochondria Transport at the Leading End

In this study, we found that RIC-7 is specifically required for mitochondria anterograde trafficking in the axon, facilitating kinesin-1-driven mitochondria movements. How does RIC-7 regulate mitochondria transport? One insight comes from its localization pattern. RIC-7 is preferentially enriched at the leading end of anterogradely trafficking mitochondria in the axon. By contrast, MIRO-1 and MTX-2 are ubiquitously distributed on mitochondria (Fu et al., 2020; Zhao et al., 2021). We also found that UNC-116, the sole ortholog of kinesin-1 in *C. elegans,* displays similar mitochondria localization as RIC-7. Previous *in vitro* studies found that kinesin-1 molecules that are artificially attached to liposomes would dynamically accumulate at the tip of an extending tube. A model was proposed in which lagging kinesin molecules will rapidly move to the growing tip, where they accumulate and exert force on the tube (Leduc et al., 2004). Our data indicate that dynamic accumulation of kinesin motors on membrane compartments occurs *in vivo*, and that RIC-7 displays similar dynamic localization.

How does RIC-7 facilitate kinesin-1 mediated mitochondria traffic? One model is that RIC-7 is an essential part of the adaptor complex. RIC-7 localization to mitochondria depends on MIRO-1 and metaxin2. From this position, RIC-7 might recruit kinesin-1 to mitochondria. As kinesin-1 accumulates at the leading end due to its motor function, RIC-7 would gather at the same spot. Indeed, when linked to kinesin-1 by a light-inducible dimerization system, Miro would accumulate at the tip of a deformed mitochondria (Song et al., 2022).

However, RIC-7 may also have a specialized function at mitochondria tips, for example, to help stabilize the clustering of kinesin-1/UNC-116. Theoretical analysis has shown that the collective force generated by accumulated kinesin-1 motors at the tip allows tube formation (Leduc et al., 2004). In the context of organelle trafficking, the accumulation of motors greatly facilitates organelle movements. For example, an *in vivo* study has demonstrated that clustering of dynein into microdomains on the membrane of a phagosome facilitates rapid directed transport of the phagosome toward microtubule minus ends. This clustering was proposed to allow cooperative force on a single microtubule (Rai et al., 2016). *In vitro*, clustering of small numbers of kinesin-1 by DNA scaffold enables long-distance transport that normally would require high numbers of individual motors (Jiang, 2021). Given the special localization and cellular function of RIC-7, RIC-7 may act as a scaffold to cluster small numbers of kinesin-1 molecules to enable their fast rebinding to microtubules and promote their processivity. Our finding that RIC-7’s function requires its disordered sequence may hint at a clustering, rather than an adaptor, role.

### Potential conservation of RIC-7 outside *C. elegans*

Previous analysis found that RIC-7 is a highly divergent protein even among nematodes, and no mammalian homologs can be found based on its primary sequence, which does not contain any known domains or motifs (Rawson et al., 2014). Using new tools for homology detection (HHPred and Dali), we found that the C-terminus of RIC-7b (471-709aa) is similar to the human Otulin domain. However, our structure/function analysis showed that this domain is dispensable for RIC-7’s function in mitochondria traffic (Fig. 8B). By contrast, we found that the 1-470aa fragment of RIC-7 is necessary and sufficient to rescue mitochondria localization defects in *ric-7* mutants. This fragment contains a relatively ordered region (1-99aa) that can localize to mitochondria, and a disordered region (100-470aa) that is required for RIC-7’s function (Fig. 8). The exact molecular function of this disordered region is unclear. Interestingly, unlike 1-470aa that mimics RIC-7’s full-length localization, 1-99aa is not confined to subdomains of mitochondria (Fig. S4). This resembles the localization of RIC-7 in *mtx-2* mutants and in kinesin-1 knockdown, suggesting that 100-470aa may involve in the interaction between RIC-7 and MTX-2 or kinesin-1. We speculate that potential functional equivalents of RIC-7 in mammalian systems may also be largely composed of a disordered region, possibly explaining why this critical factor for mitochondria traffic has evaded homology-based searches.

## Materials and Methods

### Worm Strains

All strains were maintained at 20°C on NGM plates seeded with OP50 E. coli.

**Table S1.** List of strains used in this study

| Strain | Genotype |
| --- | --- |
| XE2194 | *wpIs127[Pttx-3::mito::egfp; Pttx-3::mcherry::rab-3; Punc-122::gfp] X* |
| XE2257 | *unc-116(rh24sb79) III; wpIs127[Pttx-3::mito::egfp; Pttx-3::mcherry::rab-3; Punc-122::gfp] X* |
| XE2322 | *trak-1(wp84) I; wpIs127[Pttx-3::mito::egfp; Pttx-3::mcherry::rab-3; Punc-122::gfp] X* |
| XE3157 | *trak-1(wy50182) I; wpIs127[Pttx-3::mito::egfp; Pttx-3::mcherry::rab-3; Punc-122::gfp] X* |
| XE2357 | *miro-1(wp88) I; wpIs127[Pttx-3::mito::egfp; Pttx-3::mcherry::rab-3; Punc-122::gfp] X* |
| XE2447 | *miro-3(wp94) I; miro-1(wp88) IV; miro-2(wp92) wpIs127[Pttx-3::mito::egfp; Pttx-3::mcherry::rab-3; Punc-122::gfp] X* |
| XE2366 | *ric-7(n2657) V; wpIs127[Pttx-3::mito::egfp; Pttx-3::mcherry::rab-3; Punc-122::gfp] X* |
| XE3150 | *miro-1(wp88) IV; ric-7(n2657) V; wpIs127[Pttx-3::mito::egfp; Pttx-3::mcherry::rab-3; Punc-122::gfp] X* |
| XE2651 | *wpEx451[Pttx-3::mito::Dendra2::let-858 3'UTR+ Punc-122::gfp]* |
| XE2815 | *miro-1(wp88) IV; wpEx451[Pttx-3::mito::Dendra2::let-858 3'UTR+ Punc-122::gfp]* |
| XE3158 | *trak-1(wy50182) I; wpEx451[Pttx-3::mito::Dendra2::let-858 3'UTR+ Punc-122::gfp]* |
| XE2817 | *wpEx449[Pitr-1pB::mito::TagRFP::let-858 3' UTR + Podr-1::GFP]* |
| XE2799 | *ric-7(n2657) V; wpEx449[Pitr-1pB::mito::TagRFP::let-858 3' UTR + Podr-1::GFP]* |
| XE2798 | *ric-7(wp127[ric-7::gfp11x7]) V; shyIs36[Pmig-13::gfp1-10+Podr-1::rfp] V*<br>*wpEx483[Pitr-1pB::mito::TagRFP::let-858 3' UTR + Podr-1::GFP]* |
| XE2929 | *miro-1*(wy50180) IV; *ric-7*(wp127[*ric-7::GFP11x7*]) V; *shyIs36*[*Pmig-13::gfp1-10+Podr-1::rfp*] V; wpEx483[*Pitr-1pB::mito::TagRFP::let-858 3' UTR + Podr-1::GFP*] |
| XE2930 | *mtx-2*(wy50266) III; *ric-7*(wp127[*ric-7::gfp11x7*]) V; *shyIs36*[*Pmig-13::gfp1-10+Podr-1::rfp*] V; wpEx483[*Pitr-1pB::mito::TagRFP::let-858 3' UTR + Podr-1::GFP*] |
| XE2931 | *mtx-2*(wy50266) III; *miro-1*(wy50180) IV; *ric-7*(wp127[*ric-7::gfp11x7*]) V; *shyIs36*[*Pmig-13::gfp1-10+Podr-1::rfp*] V; wpEx483[*Pitr-1pB::mito::TagRFP::let-858 3' UTR + Podr-1::GFP*] |
| XE2917 | wpSi35[*Pitr-1 pB::ric-7b::eGFP::let-858 3'UTR + Cbun-119(+)*] I; wpEx449[*Pitr-1pB::mito::TagRFP::let-858 3' UTR + Podr-1::GFP*] |
| XE2939 | wpSi40[*Pmig-13::ric-7b::eGFP::let-858 3'UTR + Cbun-119(+)*] I; *ric-7*(n2657) V; wpEx449[*Pitr-1pB::mito::TagRFP::let-858 3' UTR + Podr-1::GFP*] |
| XE2940 | wpSi40[*Pmig-13::ric-7b::eGFP::let-858 3'UTR + Cbun-119(+)*] I; wpEx449[*Pitr-1pB::mito::TagRFP::let-858 3' UTR + Podr-1::GFP*] |
| XE3041 | wpSi43[*Pitr-1pB::ric-7b(1-470aa)::eGFP::let-858 3'UTR + Cbunc-119(+)*] I; *ric-7*(n2657) V; wpEx449[*Pitr-1pB::mito::TagRFP::let-858 3' UTR + Podr-1::GFP*] |
| XE2942 | wpSi36[*Pmig-13::ric-7b(1-470aa)::eGFP::let-858 3'UTR + Cbun-119(+)*] I; *ric-7*(n2657) V; wpEx449[*Pitr-1pB::mito::TagRFP::let-858 3' UTR + Podr-1::GFP*] |
| XE2943 | wpSi36[*Pmig-13::ric-7b(1-470aa)::eGFP::let-858 3'UTR + Cbun-119(+)*] I<br>wpEx449[*Pitr-1pB::mito::TagRFP::let-858 3' UTR + Podr-1::GFP*] |
| XE2913 | wpSi28[*Pitr-1 pB::ric-7b(1-99aa)::eGFP::let-858 3'UTR + Cbun-119(+)*] I; *ric-7*(n2657) V; wpEx449[*Pitr-1pB::mito::TagRFP::let-858 3' UTR + Podr-1::GFP*] |
| XE2933 | wpSi38[*Pmig-13::ric-7b(1-99aa)::eGFP::let-858 3'UTR + Cbun-119(+)*] I; *ric-7*(n2657) V; wpEx449[*Pitr-1pB::mito::TagRFP::let-858 3' UTR + Podr-1::GFP*] |
| XE2934 | wpSi38[*Pmig-13::ric-7b(1-99aa)::eGFP::let-858 3'UTR + Cbun-119(+)*] I;<br>wpEx449[*Pitr-1pB::mito::TagRFP::let-858 3' UTR + Podr-1::GFP*] |
| XE2885 | wpSi30[*Pitr-1 pB::ric7b(100-470aa)::eGFP::let-858 3'UTR + Cbun-119(+)*] I *ric-7*(n2657) V wpEx449[*Pitr-1pB::mito::TagRFP::let-858 3' UTR + Podr-1::GFP*] |
| XE2936 | wpSi39[*Pmig-13::ric-7b(100-470aa)::eGFP::let-858 3'UTR + Cbun-119(+)*] I; *ric-7*(n2657) V; wpEx449[*Pitr-1pB::mito::TagRFP::let-858 3' UTR + Podr-1::GFP*] |
| XE2937 | *wpSi39[Pmig-13::ric-7b(100-470aa)::eGFP::let-858 3'UTR + Cbunc-119(+)] I; wpEx449[Pitr-1pB::mito::TagRFP::let-858 3' UTR + Podr-1::GFP]* |
| XE3045 | *wpSi44[Pitr-1pB::ric-7b(471-709aa)::eGFP::let-858 3'UTR + Cbunc-119(+)] I; ric-7(n2657) V; wpEx449[Pitr-1pB::mito::TagRFP::let-858 3' UTR + Podr-1::GFP]* |
| XE2828 | *wpSi22[rab-3p::ric-7a + Cbunc-119(+)] I; ric-7(n2657) V; wpIs127[Pttx-3::mito::egfp; Pttx-3::mcherry::rab-3; Punc-122::gfp] X* |
| XE2829 | *wpSi23[rab-3p::human otulin + Cbunc-119(+)] I; ric-7(n2657) V; wpIs127[Pttx-3::mito::egfp; Pttx-3::mcherry::rab-3; Punc-122::gfp] X* |
| XE2830 | *wpSi24[rab-3p::ric-7b 471-709aa+ Cbunc-119(+)] I; ric-7(n2657) V; wpIs127[Pttx-3::mito::egfp; Pttx-3::mcherry::rab-3; Punc-122::gfp] X* |
| XE2831 | *wpSi25[rab-3p::human otulinL + Cbunc-119(+)] I; ric-7(n2657) V; wpIs127[Pttx-3::mito::egfp; Pttx-3::mcherry::rab-3; Punc-122::gfp] X* |
| XE3000 | *unc-116(wp160[unc-116::3xgfp11]) III; shyIs36[Pmig-13::gfp1-10+Podr-1::rfp] V* |
| XE3005 | *unc-116(wp160[unc-116::3xgfp11]) III; shyIs36[Pmig-13::gfp1-10+Podr-1::rfp] V; wpEx449[Pitr-1pB::mito::TagRFP::let-858 3' UTR + Podr-1::GFP]* |
| XE3134 | *shyIs53[mig-13p::AtTir1+odr-1p::gfp] II; unc-116(shy92[unc-116::AID]) III; ric-7(wp127[ric-7::GFP11x7]) shyIs36[Pmig-13::gfp1-10+Podr-1::rfp] V; wpEx527[Pitr1-pB::mito::TagRFP::let858 3'UTR + Pmyo-2::mCherry]* |

### Mitochondria marker selection

In initial attempts to label mitochondria in AIY, we tried three different markers: TOMM-20 transmembrane domain, cytochrome c oxidase subunit 4 (COX-4), and the mitochondria matrix localization signal (MLS) from the chicken mitochondrial aspartate aminotransferase as used in pPD96.32 (Fire Lab Vector Kit). Overexpression of TOMM-20::eGFP exhibited inconsistent mitochondria morphology, sometimes tubular and sometimes globular. Overexpression of COX-4::eGFP resulted in mitochondria fragmentation. In contrast, mitochondria labeled by MLS::eGFP (mito::eGFP in this paper) show consistently tubular morphology. Therefore, we decided to pursue with this marker. The microinjection mix contained 30 ng/μL Pttx-3::mito::egfp::let-858 3’UTR, 30 ng/ul DACR18 (Pttx-3::mcherry::rab-3::unc-10 3’UTR), 25 ng/ul Punc-122::GFP and Promega 1 kb DNA ladder as the filler. The mix was injected into N2, and transgenic animals with integrated extrachromosomal arrays were then generated by the TMP/UV method. Among several integration lines, we selected *wpIs127* because it expresses the transgenes at a relatively low level and exhibits consistent mitochondria morphology. *wpIs127* was outcrossed five times before further analysis.

Mitochondria in DA9 were labeled by mitochondria matrix localization signal (pPD96.32, Fire Lab Vector Kit) fused with TagRFP and expressed as extrachromosomal arrays. The microinjection mix contained Pitr-1pB::mito::TagRFP::let858 3’UTR at 30 ng/μL, Podr-1::gfp at 20 ng/μL and Promega 1 kb DNA ladder as the filler.

### Molecular Cloning

All plasmids used for expressing transgenes in neurons were assembled to the destination vector pCFJ150 using Gateway recombination (Invitrogen). Gibson reactions were used to generate the entry clones (Gibson et al., 2009). Plasmids that express *trak-1* and *miro-1* sgRNAs were made by swapping the N_19_ sequence of the *unc-119* sgRNA from the preexisting plasmid (Addgene #46169) (Friedland et al., 2013) using PCR, followed by 5’ phosphorylation and blunt end ligation.

**Table S2.**
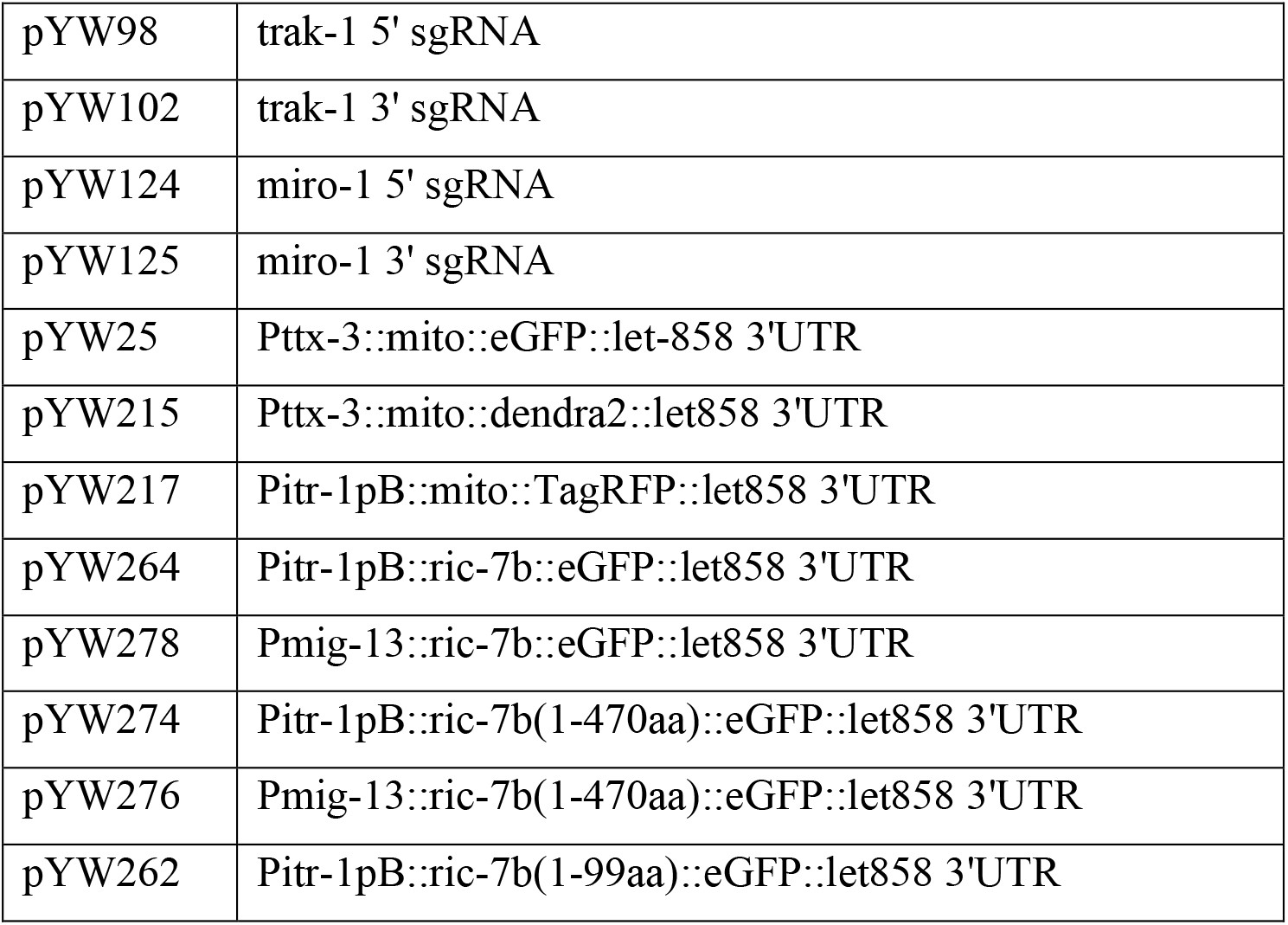
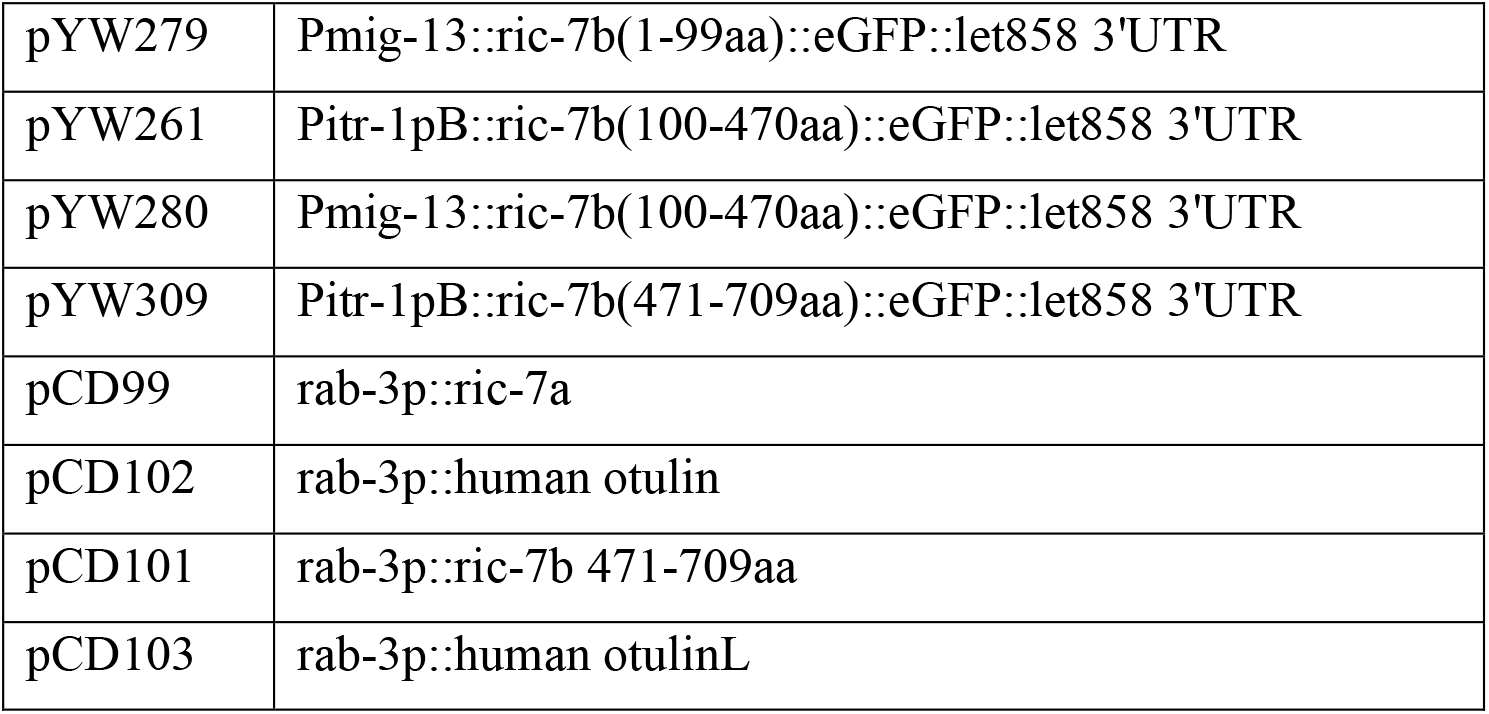
List of plasmids used in this study:

|  |  |
| --- | --- |
| pYW98 | trak-1 5' sgRNA |
| pYW102 | trak-1 3' sgRNA |
| pYW124 | miro-1 5' sgRNA |
| pYW125 | miro-1 3' sgRNA |
| pYW25 | Pttx-3::mito::eGFP::let-858 3'UTR |
| pYW215 | Pttx-3::mito::dendra2::let858 3'UTR |
| pYW217 | Pitr-1pB::mito::TagRFP::let858 3'UTR |
| pYW264 | Pitr-1pB::ric-7b::eGFP::let858 3'UTR |
| pYW278 | Pmig-13::ric-7b::eGFP::let858 3'UTR |
| pYW274 | Pitr-1pB::ric-7b(1-470aa)::eGFP::let858 3'UTR |
| pYW276 | Pmig-13::ric-7b(1-470aa)::eGFP::let858 3'UTR |
| pYW262 | Pitr-1pB::ric-7b(1-99aa)::eGFP::let858 3'UTR |
| pYW279 | Pmig-13::ric-7b(1-99aa)::eGFP::let858 3'UTR |
| pYW261 | Pitr-1pB::ric-7b(100-470aa)::eGFP::let858 3'UTR |
| pYW280 | Pmig-13::ric-7b(100-470aa)::eGFP::let858 3'UTR |
| pYW309 | Pitr-1pB::ric-7b(471-709aa)::eGFP::let858 3'UTR |
| pCD99 | rab-3p::ric-7a |
| pCD102 | rab-3p::human otulin |
| pCD101 | rab-3p::ric-7b 471-709aa |
| pCD103 | rab-3p::human otulinL |

### Generation of Knock-Outs and Knock-Ins with CRISPR/Cas9

*trak-1(wp84)*, *eat-3(wp85)*, *miro-1(wp88)*, *miro-2(wp92)* and *miro-3(wp94)* are whole gene deletions generated by NHEJ with CRISPR/Cas9. For *trak-1(wp84)* and *miro-1(wp88)*, CRISPR/Cas9 was performed using the *dpy-10(cn64)* as the co-conversion marker as previously described in Arribere et al., 2014. Briefly, sgRNA and Cas9 was expressed using plasmids. The microinjection mix contained 50 ng/μl sgRNA vector, 40 ng/μl dpy-10 sgRNA vector, 50 ng/μl Cas9 expression vector (pDD162) (Dickinson et al., 2013), and 26 ng/ul dpy-10 repair template AF-ZF-827 (Arribere et al., 2014). The mix was injected into XE2194, and F1 rollers were genotyped by single worm PCR. *miro-2(wp92)* and *miro-3(wp94)* were generated by NHEJ using preassembled Cas9 ribonucleoprotein complex as described in Dokshin et al., 2018. The injection mix contains 0.25 μg/μl S.pyogenes Cas9 (IDT# 1081058), 0.1 μg/μl tracrRNA (IDT# 1072532), 0.056 μg/μl crRNA in total (0.028 μg/μl for each sequence, synthesized by IDT), and 40ng/μl PRF4 (rol-6 (su1006)) plasmid. *miro-2* and *miro-3* were simultaneously targeted by the same sgRNAs, and the mix was injected into *miro-1(wp88)* to generate *miro* triple mutant. F1 progenies (rollers and non-rollers) were screened by single worm PCR.

*ric-7::7xgfp11* knock-in was generated by CRISPR/cas9 using preassembled Cas9 ribonucleoprotein complex as previously described (Ghanta and Mello, 2020). Briefly, the injection mix contained 0.25 μg/μl S.pyogenes Cas9 (IDT# 1081058), 0.1 μg/μl tracrRNA (IDT# 1072532), 0.056 μg/μl crRNA (synthesized by IDT), 25ng/μl pre-melted double-stranded DNA repair template or 0.11 μg/μl of ssDNA repair template and 40ng/μl PRF4 (rol-6 (su1006)) plasmid. Non-roller F1 progeny from P0 roller jackpot plates were screened by single worm PCR. Homozygotes from the F2 generation were verified by Sanger sequencing.

For *unc-116::3xgfp11* and *unc-116::AID* knock-in, instead of using tracrRNA and crRNA, a single gRNA of 3.7 μg was used. This gRNA was synthesized from ssDNA using the EnGen sgRNA Synthesis Kit, S. pygenes (NEB# E3322S) and subsequently purified using a Monarch RNA cleanup kit (NEB#T2040).

All repair templates contained 35 bp homology arms on both ends. For tandem *gfp11* insertions, a GSGGGG liker was added between the gene of interest and *gfp11*. For *unc-116*, we chose to use 3x tandem repeats instead of 7x, as mitochondria trafficking was dramatically reduced when *unc-116* was tagged with 7xgfp11. The repair templates for *ric-7::7xgfp11* and *unc-116::3xgfp11* were synthesized as a Gblock by IDT, amplified by PCR, gel purified and further cleaned up with the Qiagen MinElute kit. The repair template for *unc-116::AID* was synthesized by IDT as ssDNA.

#### *ric-7::7xgfp11* repair template

GTCCGGTTCCCCTGTTCTACATACCGGACGATGAAggatccggaggtggcgggCGTGACCACATGGTCCTTCATGAGTATGTAAATGCTGCTGGGATTACAGGTGGCTCTGGAGGTAGAGATCATATGGTTCTCCACGAATACGTTAACGCCGCAGGCATCACTGGCGGTAGTGGAGGACGCGACCATATGGTACTACATGAATATGTCAATGCAGCCGGAATAACCGGAGGGTCCGGAGGCCGGGATCACATGGTGCTGCATGAGTATGTGAACGCGGCGGGTATAACTGGTGGGTCGGGCGGACGAGACCATATGGTGCTTCACGAATACGTAAACGCAGCTGGCATTACTGGCGGATCAGGTGGCAGGGATCACATGGTACTCCATGAGTACGTGAACGCTGCTGGAATCACAGGCGGTAGCGGCGGTCGGGACCATATGGTCCTGCACGAATATGTCAATGCTGCCGGTATCACCTAAtaactaggaccctctttttgttattcgtaacttgtaa

#### *unc-116::3xgfp11* repair template

GACAAGTGTATACGTCTCCGTCAGCAGGAATGTCAggatccggaggtggcgggCGTGACCACATGGTCCTTCATGAGTATGTAAATGCTGCTGGGATTACAGGTGGCTCTGGAGGTAGAGATCATATGGTTCTCCACGAATACGTTAACGCCGCAGGCATCACAGGCGGTAGCGG CGGTCGGGACCATATGGTCCTGCACGAATATGTCAATGCTGCCGGTATCACCCAAGGAGCTCCAAATGGCTCAAACGgtgtgtttag

#### *unc-116::AID* repair template

GACAAGTGTATACGTCTCCGTCAGCAGGAATGTCAATGCCTAAAGATCCAGCCAAACCTCCGGCCAAGGCACAAGTTGTGGGATGGCCACCGGTGAGATCATACCGGAAGAACGTGATGGTTTCCTGCCAAAAATCAAGCGGTGGCCCGGAGGCGGCGGCGTTCGTGAAGCAAGGAGCTCCAAATGGCTCAAACGgtgtgtttag

**Table S3.**
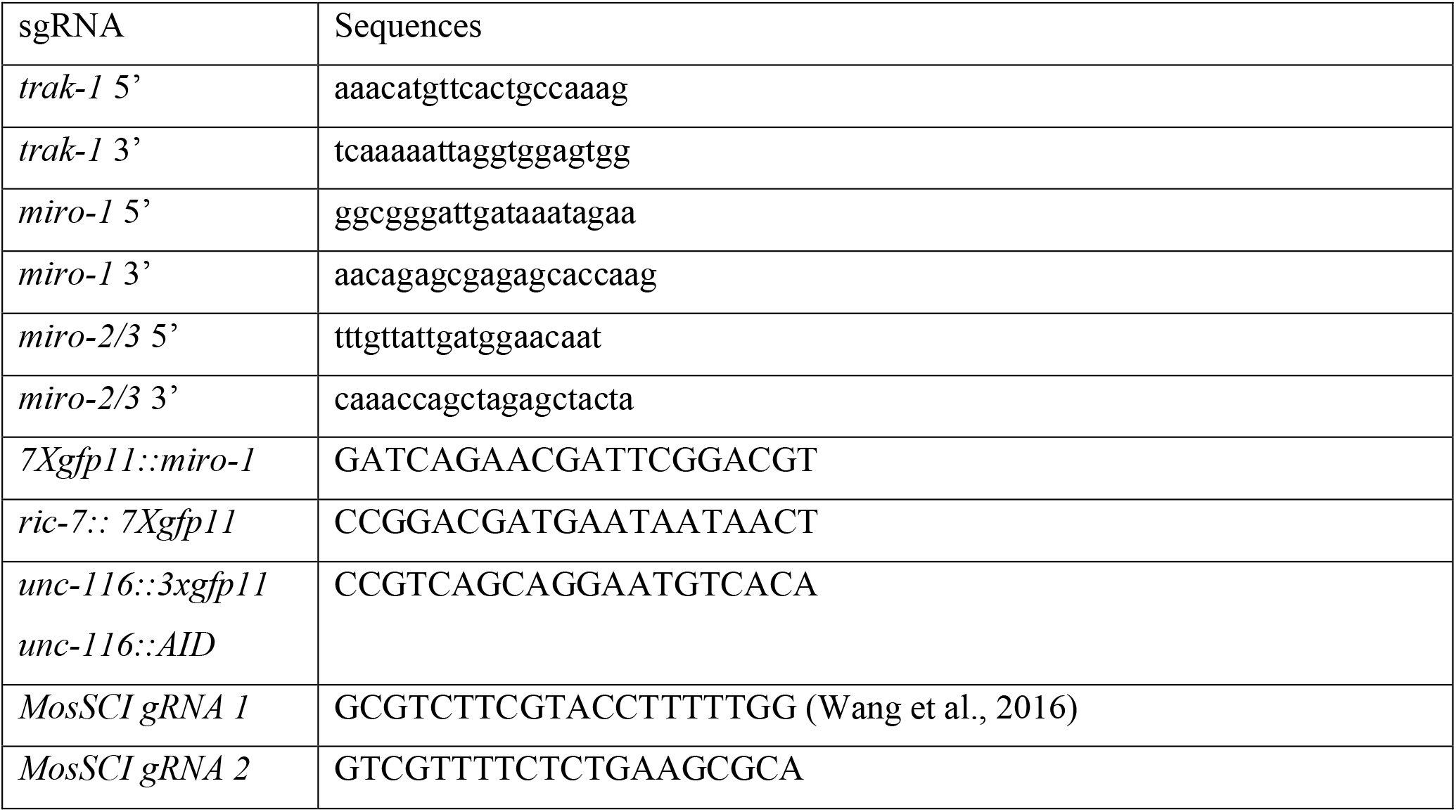
List of gRNA sequences used in this study

### Structure/function analysis of RIC-7

Structure/function analysis of RIC-7 was performed by expressing different transgenes as single copy insertions at the same genomic locus under the same promoter to achieve comparable levels of expression. As the endogenous expression level of RIC-7 in DA9 is extremely low (Fig. S5A) (Taylor et al., 2021), we initially used a weak promoter, *Pitr-1pB*, to drive the expression of transgenes in DA9 (Klassen and Shen, 2007; Taylor et al., 2021). Using this approach, mitochondria localization is restored when *ric-7b* full length is expressed, suggesting that the expression level is high enough for the functional assessment of the truncated proteins. However, it is not high enough for us to reliably examine the localization of the proteins. Therefore, we switched to a stronger DA9 promoter, *Pmig-13* (Klassen and Shen, 2007) (Fig. S5A), to examine protein localization.

CRISPR/Cas9 was used to generate single copy insertion at the ttTi5605 landing site on Chromosome I in the strain EG8078. The injection mix contained 0.25 μg/μl S.pyogenes Cas9 (IDT# 1081058), 0.1 μg/μl tracrRNA (IDT# 1072532), 0.028 μg/μl crRNA each (synthesized by IDT), the transgene plasmid constructed in pCFJ150 (100 ng/μl), pMA122 (peel-1 negative selection, 10ng/μl) and pCFJ90 (Pmyo2::mCherry, 1.5 ng/μl). Each injected P0 animal was singled and grew for 7-8 days at room temperature until starvation followed by heat shock at 34°C for 4h. The next day after heat shock, P0 plates were checked for potential insertions. Successful insertions should survive the heat shock, not express mCherry and be non-unc. ∼10 non-unc worms from each P0 plate were singled and verified for insertion by genotyping.

### Auxin Induced Single Cell Specific Knock Down Of *kinesin-1*/*unc-116*

To acutely knock down *kinesin-1*/*unc-116* in DA9, young L4 animals were cultured on NGM plates containing 1 mM Auxin (Thermo-scientific, #A10556.14) for 5 hours before imaging. Animals of the same stage cultured on NGM plates lacking auxin were served as the controls.

### Microscopy

To visualize mitochondria distribution in AIY and DA9, L4 animals were immobilized by 8 mM levamisole and mounted on 3% agarose pad. Still images were acquired with an ORCA-Flash4.0 camera (Hamamatsu) on a VT-iSIM system (BioVision) built around a Leica Laser Safe DMi8 Inverted scope with a PL APO 100x/1.47 OIL CORR TIRF objective for AIY, or a HC PL APO 63x/1.40 OIL CS2 objective for DA9. Image acquisition was controlled by MetaMorph Advanced Confocal Acquisition Software Package. The AIY neuron that is closer to the coverslip was imaged. Z-stacks were acquired with a step size of 0.2 μm to capture the whole neuron.

For time lapse imaging to visualize mitochondria trafficking in AIY, young L4 animals were immobilized by 8 mM levamisole for 5 minutes, and then washed with M9 for at least 5 times before mounting on 3% agarose pad. Movies were taken under the VT-iSIM system with 0.2 μm z-step size at 1 min interval.

To better image the endogenous proteins and mitochondria in DA9, L4 animals were immobilized with 50 mM Muscimol for 2-3 min before mounting, which increases the chance that either the dorsal or the ventral side faces the coverslip. Images were acquired with a HC PL APO 63x/1.40 OIL CS2 objective. Emission filters 525/50nm and 605/52nm were used for 488nm and 561nm illumination, respectively. Images were taken at 400ms exposure sequentially for 488nm and 561nm at each Z position. Only the dendrite or the axon was imaged for each animal, depending on the orientation. Time lapse imaging were performed for 10 min with 10-second intervals, with three z-steps of 0.2 μm step size.

FRAP of RIC-7::7xSplitGFP was performed on a Nikon Ti-2E inverted microscope system equipped with Yokogawa CSU-W1 SoRa dual disk confocal with photostimulation scanner module. 525/36nm and 605/52nm filters were used to visualize GFP and TagRFP, respectively. Images were taken with a Nikon CFI SR HP Plan Apo lambda S 100XC Sil objective using Hamamatsu ORCA-Fusion BT sCMOS camera, with 3 images of 2s intervals before photobleaching, and 30 images of 10s intervals after photobleaching for 5 min. Acquisition was driven by NIS-Elements software.

To generate the kymograph of kinesin-1::3xsplitGFP (UNC-116::3xsplitGFP) trafficking, images were taken continuously on a single plane at 400ms exposure with the 488nm laser using the streaming mode.

For Photoconversion, images were acquired on a UltraVIEW VoX spinning-disc confocal microscope (PerkinElmer) built around a Nikon Ti-E Eclipse equipped with a 60x/1.4NA CFI Plan Apo objective and a Hamamatsu C9100-50 camera and driven by the Volocity software (Improvision). Mitochondria in the cell body or in the axon in L2 worms were photoconverted with 405nm laser using the PhotoKinesis Unit. Worms were recovered and placed on NGM plates with OP50. 24 hours later, these worms were imaged again.

### Image analysis

To quantify mitochondria in the AIY axon, we used two different metrics: 1) the fraction of the total fluorescence of mito::eGFP in the axon; 2) Number of mitochondria in the axon. We arrived at the same conclusion using both metric. The first metric is defined by the following formula: the total intensity of the axon/(the total intensity of the cell body + the axon), which was calculated with a custom Matlab script, using maximum projection images. To quantify mitochondria in DA9, only the 2^nd^ metric was used.

To quantify kinesin-1::3xsplitGFP (UNC-116::3xsplitGFP) enrichment at the leading end of anterogradely trafficking mitochondria, linescans of the GFP fluorescence intensity along the length of mitochondria were generated in FIJI. After background (intensities outside of the neuron but inside the worm) subtraction, the intensity values were averaged across different timepoints over the course of the movements, including the period when mitochondria were paused, and then normalized to the maximum value of all timepoint. To average across different mitochondria that differ in length, the values at 51 regularly spaced points along the mitochondria length were estimated using linear interpolation with the interp1 function in Matlab.

Kymographs of kinesin-1::3xsplitGFP (UNC-116::3xsplitGFP) trafficking were generated and analyzed using KymoToolBox.

To compare RIC-7 localization in different genotypes, two different methods were used: 1) Enrichment on Mitochondria (Figure 7D); 2) Pearson’s Correlation (Figure 7E). Enrichment on Mitochondria is defined as the ratio between the intensity of RIC-7::7xSplitGFP puncta on mitochondria and the cytosolic GFP signals, both of which were manually measured in ImageJ and background subtracted. For *mtx-2* and *miro-1; mtx-2* double mutants where RIC-7::7xSplitGFP does not form discrete puncta, the GFP intensity on the entire mitochondria was measured. For Pearson’s Correlation, the DA9 cell body excluding the nucleus was manually specified in ImageJ, and then a customized macro in ImageJ was used to get the intensities of each pixel in both channels. Pearson’s Correlation between RIC-7::7xSplitGFP and mito::TagRFP was calculated using the Matlab function corrcoef.

### Statistical Analysis

All statistical analysis was performed in GraphPad Prism. Two-sided Fisher’s exact tests were used for binary data. For non-binary data, t-test was used to compare two groups, and one-way ANOVA test followed by Dunn’s multiple comparisons test was used to compare multiple groups.

### Data Availability Statement

All data are available in the main text or the supplementary materials.

## Acknowledgements

We would like to thank the International Caenorhabditis Genetics Center for providing *C. elegans* strains, members in the Hammarlund Lab and Yogev Lab for technical support and discussion, Shawn Ferguson for suggestions on structure/function analysis. We thank Dr. Erik E. Griffin, Dr. Jun Hyun Park, Dr. Lauren Panzera, Hadas Dabas for providing feedback on the manuscript. The work is supported by NIH (R01 NS094219).

The authors declare no competing financial interests.

## Author contributions

Author contributions: Y. Wu and M. Hammarlund conceived the study. Y. Wu performed and analyzed most of the experiments. C. Ding contributed to the structure/function analysis. A. Weinreb performed the analysis of *ric-7* splicing variants and helped with image analysis. L. Mannings and D. A. Colon-Ramos performed the EM reconstruction of AIY neurons. G. Swaim and S. Yogev created the *unc-116::AID* strain. The manuscript was written by Y. Wu and M. Hammarlund, with input from all co-authors.

---

**Figure S1.**
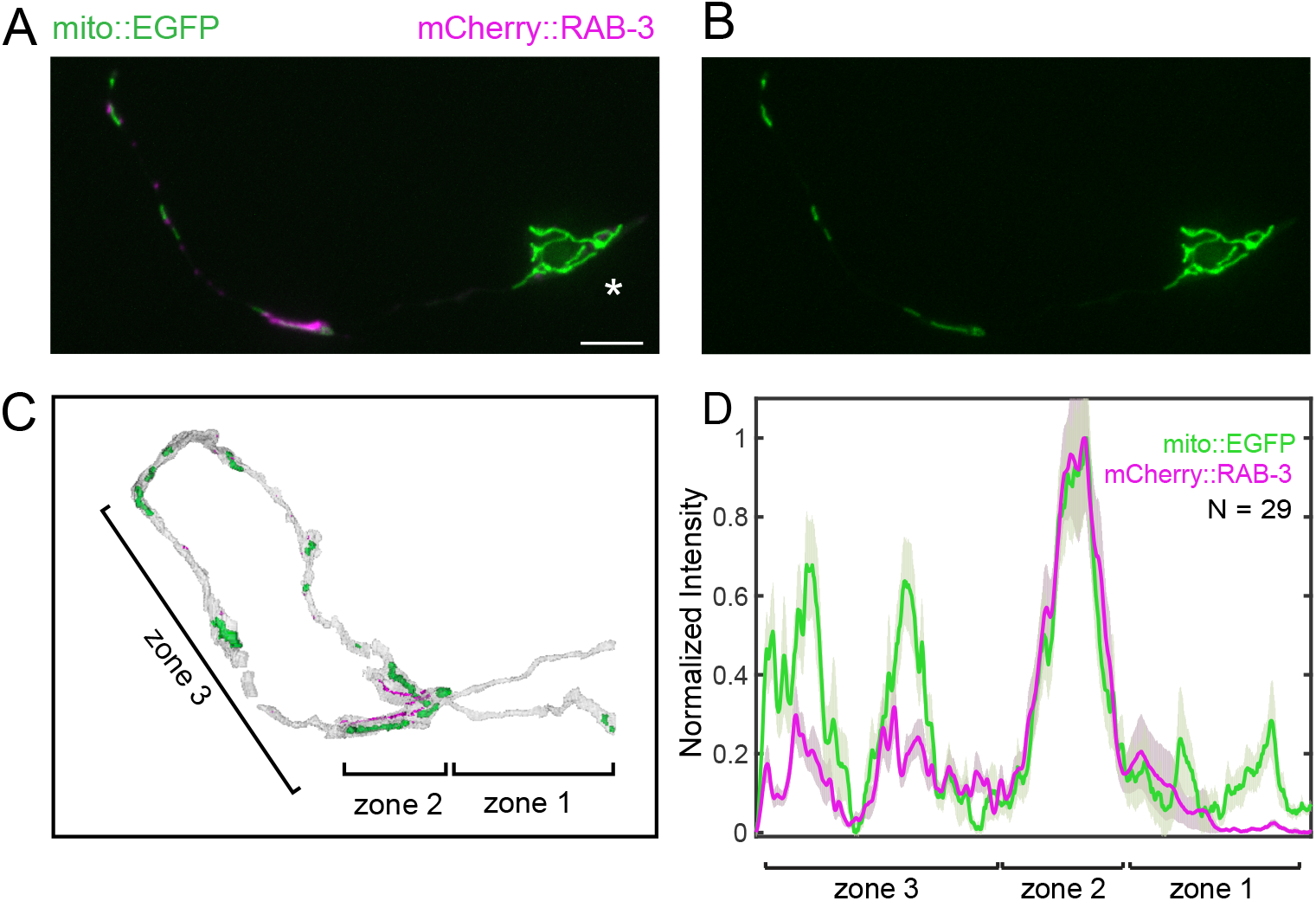
Validation of a matrix-targeted eGFP mitochondria marker in the AIY neuron. **(A-B)** Representative images of mitochondria labeled by the matrix-targeted EGFP and synaptic vesicles labeled by mCherry::RAB-3 in one AIY neuron, showing similar distribution as in the EM reconstruction. Zone 2 mitochondria often exhibit elongated morphology, as in the EM reconstruction. Asterisk indicates the cell body. Scale bar = 5 μm. **(C)** EM construction of the two AIY neurites with mitochondria and dense projections (representing synapses) highlighted in green and magenta, respectively (White et al., 1986). Mitochondria appear to accumulate at zone 2 and two other places in zone 3. **(D)** The averaged line scans of mito::eGFP and mCherry::RAB-3 in the AIY neurites. Mitochondria are enriched at zone 2 and two other places in zone 3.

**Figure S2.**
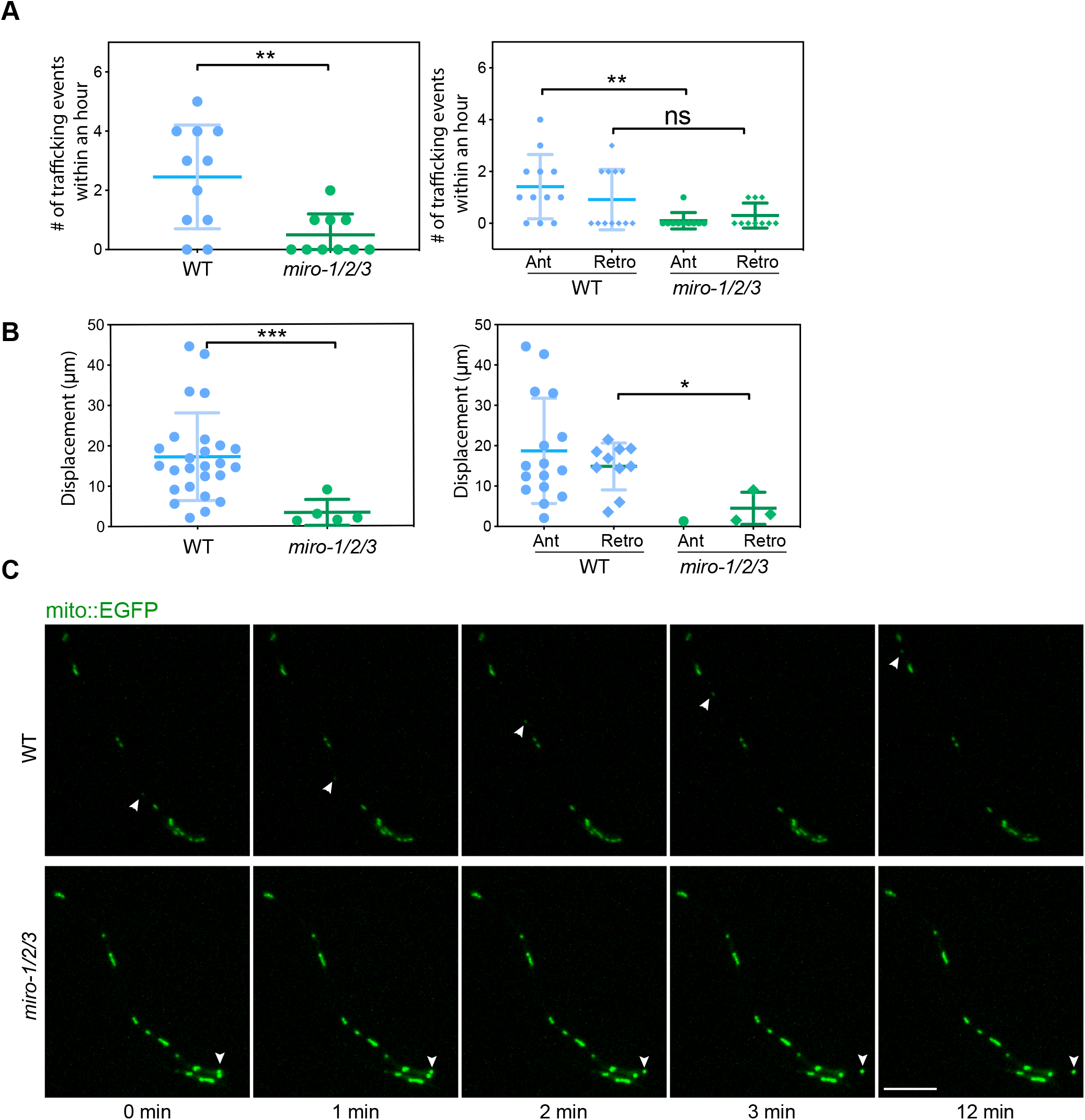
Mitochondria Trafficking is reduced in *miro-1/2/3* mutant. **(A)** Number of trafficking events within an hour in AIY axon. Left panel: combining both anterograde and retrograde; Right panel: same dataset as in the left panel (1 oscilatory event of both genotypes was excluded from this graph). ** p < 0.01, ns = not significant (Mann-Whitney test) **(B)** Displacement of all trafficking events observed. Left panel: combining both anterograde and retrograde; Right panel: same dataset as in the left panel (1 oscilatory event of both genotypes was excluded from this graph). No statistical test was performed between WT anterograde and *miro-1/2/3* anterograde as there is only 1 datapoint for the latter. * p < 0.05, *** p < 0.001 (t-test) **(C)** Representative images showing a trafficking event in WT and *miro-1/2/3* mutant, highlighted by the arrow heads. Note the short displacement of the mitochondrion in *miro-1/2/3* mutant. Cell body is to the right, which is cropped. Scale bar = 5 μm.

**Figure S3.**
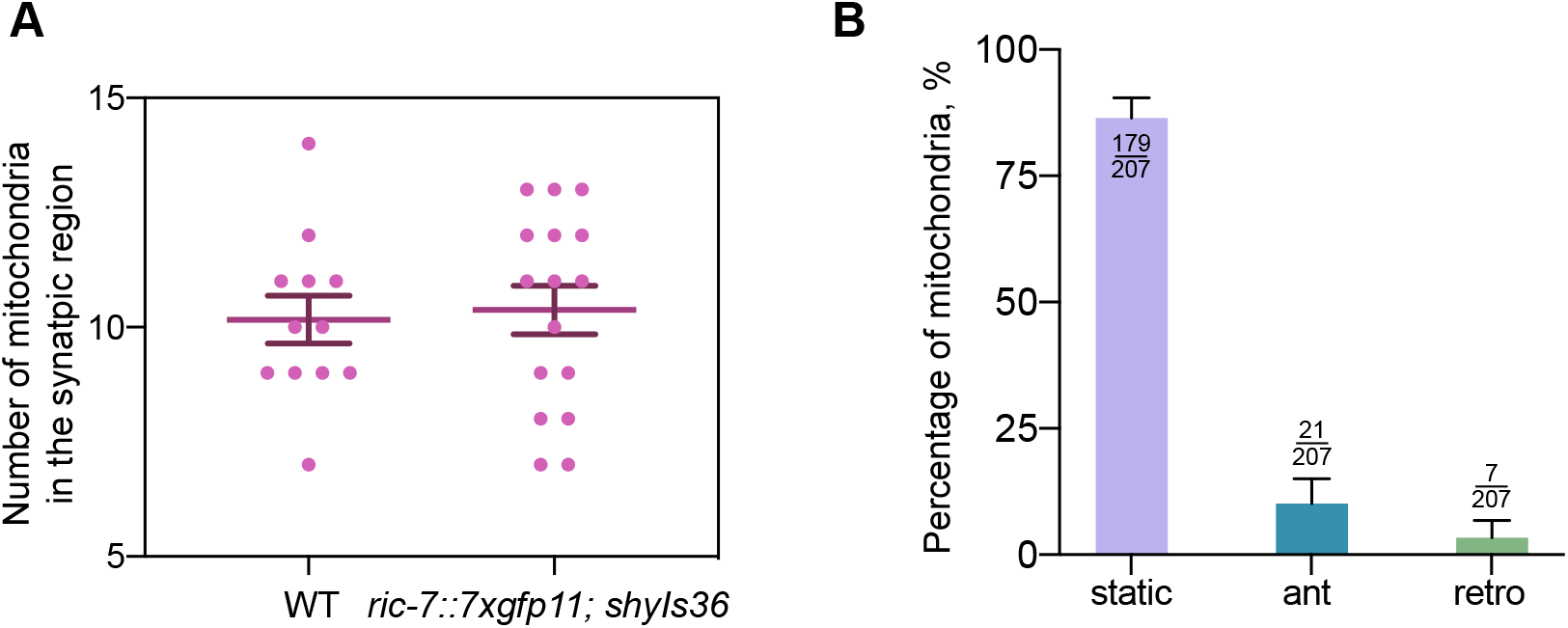
Functional validation of RIC-7::7xsplitGFP. **(A)** Mitochondria distribution in the axon is not affected when RIC-7 is tagged with 7xsplitGFP at its C-terminus. WT data is duplicated from Fig. 3E. Error bars represent SEM. **(B)** Percentage of mobile mitochondria in the DA9 axon. Events in the synaptic region are quantified and the numbers are indicated at the top of each bar. Error bars represent 95% confidence intervals.

**Figure S4.**
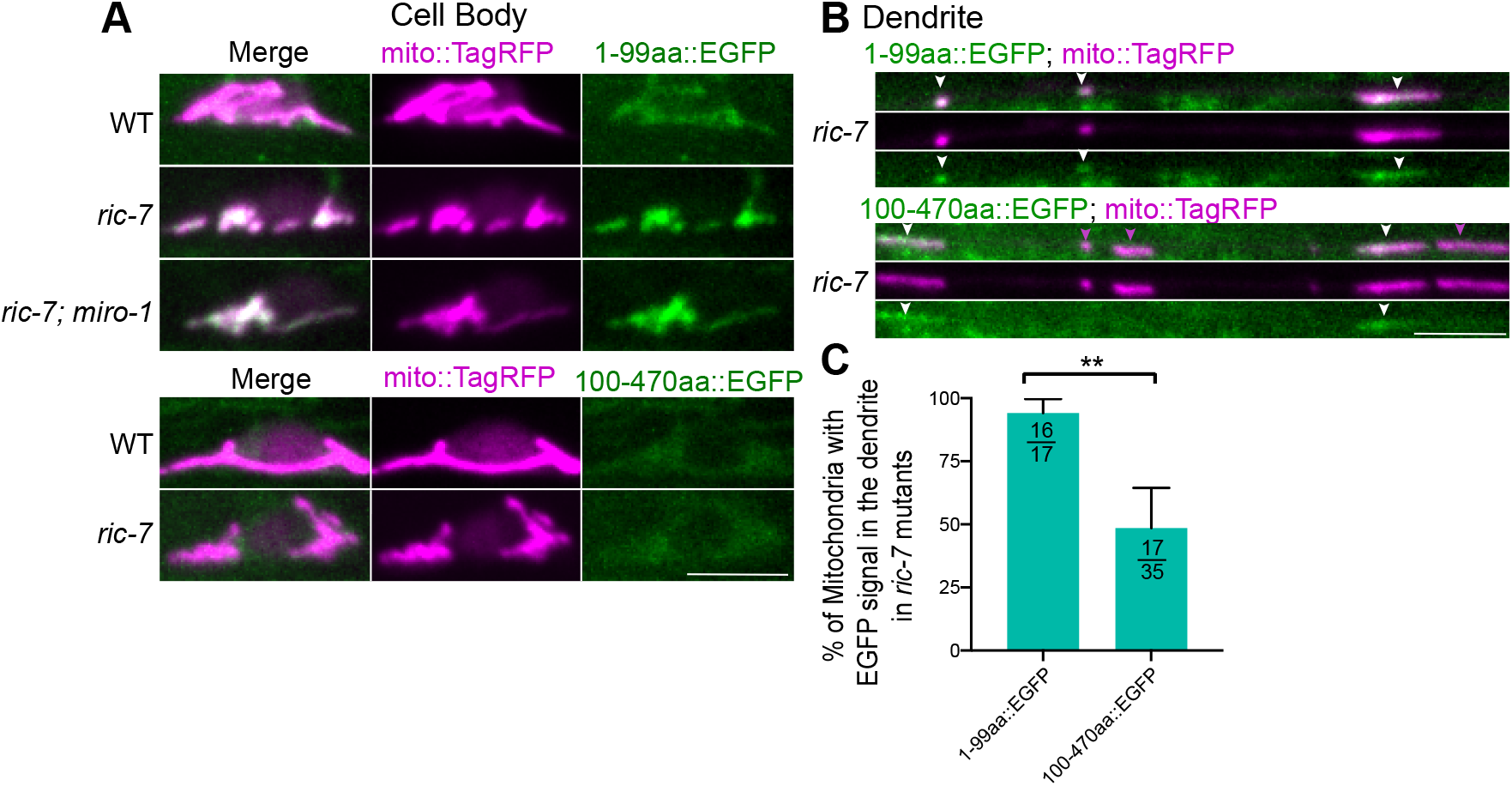
Localization of RIC-7b(1-99aa) and RIC-7b(100-470aa) in the cell body and in the dendrite expressed as transgenes driven by the promoter of *Pmig-13* (see methods). **(A)** Representative images showing the localization of 1-99aa and 100-470aa in the cell body in different genotypes. **(B)** Representative images showing the localization of 1-99aa and 100-470aa in the dendrite in *ric-7* mutant animals. **(C)** Quantification of the percentage of mitochondria associated with EGFP signals. Number of mitochondria counted are indicated at the top of each bar. Error bars represent 95% confidence intervals. ** p < 0.01.

**Figure S5.**
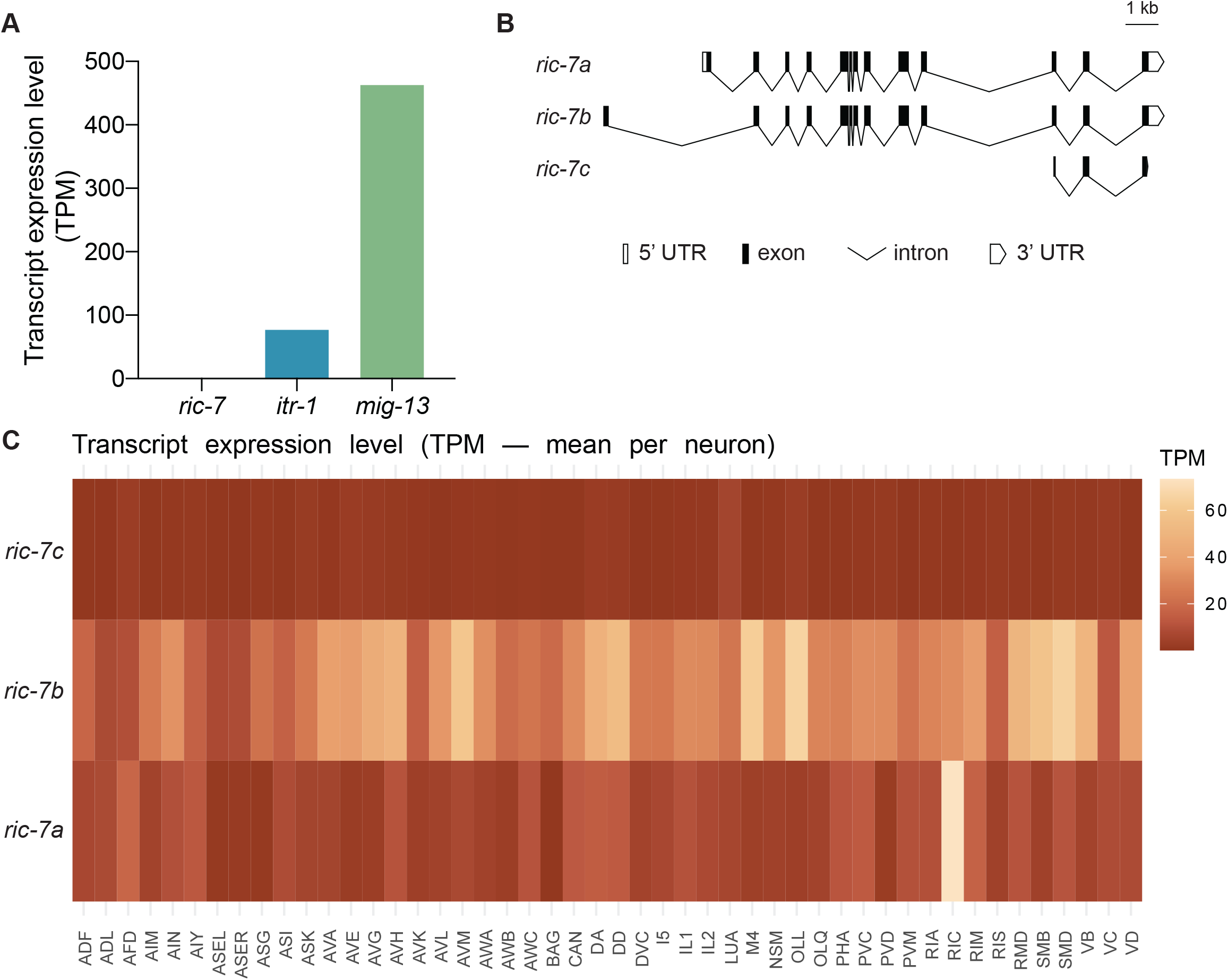
Transcript expression level of *ric-7, itr-1 and mig-13* from the CeNGEN dataset. (A) Transcript expression level of *ric-7, itr-1 and mig-13* in DA9 (Threshold=2) (Taylor et al., 2021). *ric-7* is so lowly expressed that it’s not detected in DA9. (B) Schematic representations of the three splicing isoforms of *ric-7*. (C) Transcript expression level of the three splicing isoforms of *ric-7* in different neurons. Note that *ric-7b* has a relatively higher expression compared with the other two isoforms.

**Figure S6.**
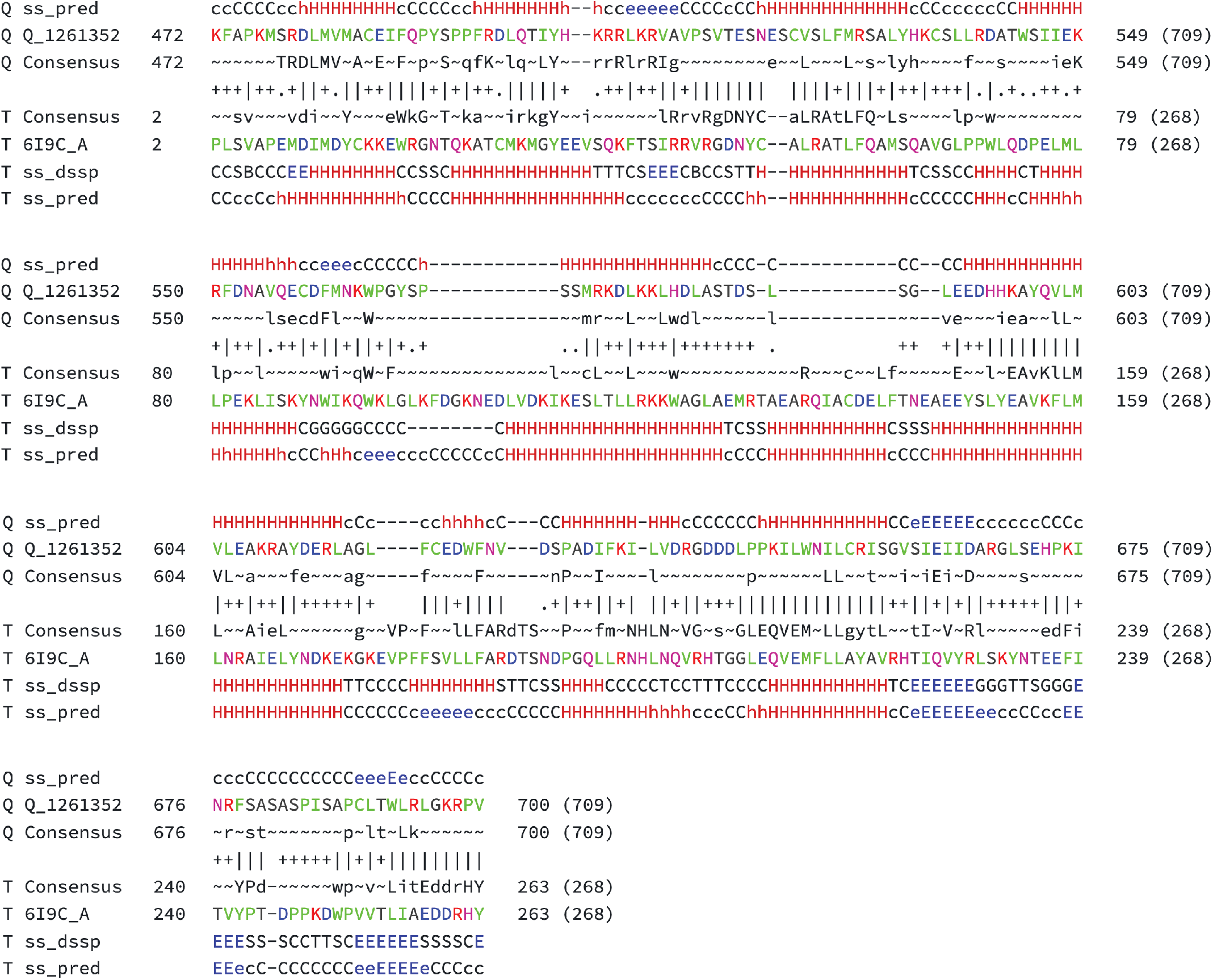
Protein remote homology detection by HHPred. Protein remote homology detection for RIC-7b was performed using HHPred (https://toolkit.tuebingen.mpg.de/tools/hhpred). Human proteome was selected as the search database. The top two hits were OtulinL and Otulin. Sequence alignment between RIC-7b and Otulin is shown in this figure. Note the high similarity of the secondary structures of the Query (Q1261352: RIC-7b) and Target (T 619C_A: Otulin). SS_Pred denote the PSI-PRED secondary structure prediction and ‘ss_dssp’ is the actual secondary structure determined by the program DSSP. Quality of the column–column match: ‘|’ very good, ‘+’ good, ‘·’ neutral, ‘−’ bad and ‘=’ very bad.

**Figure S7.**
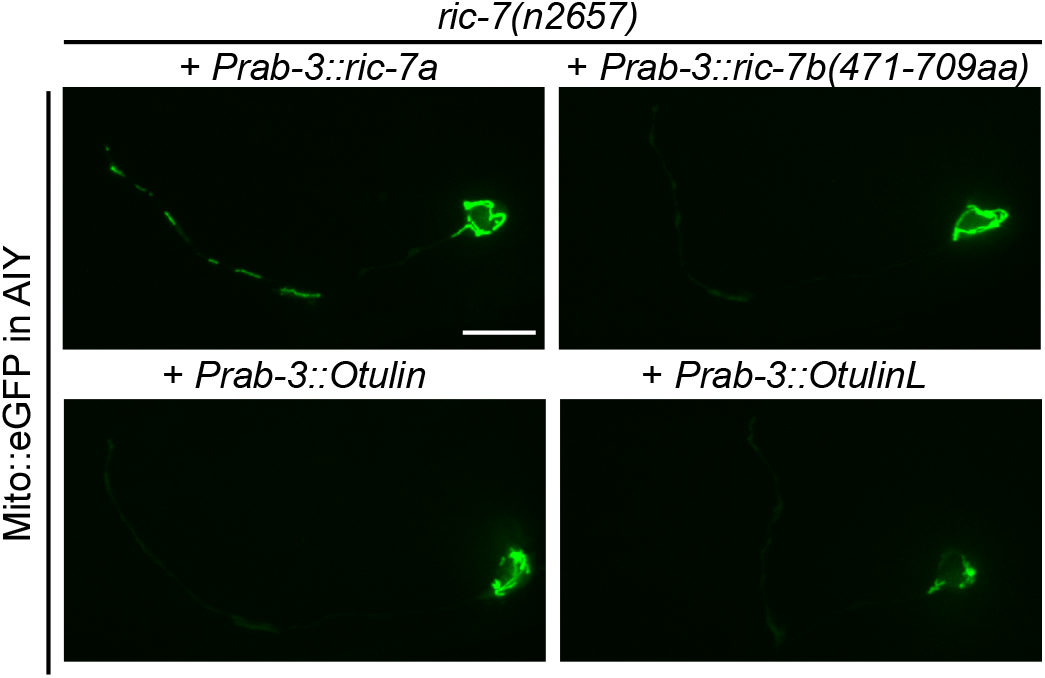
Human Otulin or OtulinL, or the C-term domain of RIC-7 cannot rescue mitochondria localization in *ric-7* mutant. Representative images showing mitochondria localization in *ric-7* mutant animals expressing the human Otulin or OtulinL, or the C-term domain (471-709aa) of RIC-7 driven by a pan-neuronal promoter from a single copy insertion.

**Movie S1: Mitochondria trafficking in AIY neuron in WT animals**

Representative movie of mitochondria trafficking in WT animals. One of the AIY neurons expressing mito::EGFP was imaged once every minute for one hour under the iSIM. The display rate is 5fps. A few trafficking events (dimmer mitochondria) were observed within an hour. See Fig. S2 C for the still images.

**Movie S2: Mitochondria trafficking in AIY neuron in *miro-1/2/3* mutant animals** Representative movie of mitochondria trafficking in *miro-1/2/3* mutant animals. One of the AIY neurons expressing mito::EGFP was imaged once every minute for one hour under the iSIM. The display rate is 5fps. Only 1 trafficking event with a short distance was observed. See Fig. S2 C for the still images.

**Movie S3: RIC-7::7xsplitGFP signals are associated with anterograde but not retrograde trafficking mitochondria**

Ventral view of an axon segment that is proximal to the cell body. RIC-7::7xsplitGFP in green and mito::TagRFP in magenta. Circle indicates the cell body that is out of focus, with the axon to the right. One mitochondrion is transported back to the cell body without RIC-7::7xsplitGFP, whereas another mitochondrion transporting from the cell body into the axon is associated with RIC-7::7xsplitGFP. Images were taken every 10s and play rate is 5fps.

**Movie S4: RIC-7::7xsplitGFP are enriched at the leading end of mitochondria that transported anterogradely in the axon**

Dorsal view of the synaptic region in the axon. Distal end is to the left. RIC-7::7xsplitGFP in green and mito::TagRFP in magenta. A mitochondrion is moving anterogradely, with RIC-7::7xsplitGFP at the leading end. Images were taken every 10s and play rate is 5fps. See Fig. 4I for the still images.

**Movie S5: RIC-7::7xsplitGFP are enriched at the leading end of mitochondria that enter the axon**

Ventral view of the cell body with the axon to the right. The dendrite is out-of-focus. RIC-7::7xsplitGFP in green and mito::TagRFP in magenta. This movie shows two mitochondria that first extend into the axon, followed by fission and transport, with RIC-7::7xsplitGFP at the leading end. Images were taken every 10s and play rate is 5fps. See Fig. 4K for the still images. Images were taken every 10s and play rate is 5fps.

**Movie S6: RIC-7::7xsplitGFP are not associated with mitochondria that enter the dendrite** Ventral view of the cell body with the dendrite to the right and the axon to the left. The axon is slightly out of focus. RIC-7::7xsplitGFP in green and mito::TagRFP in magenta. One mitochondrion enters the dendrite without RIC-7::7xsplitGFP. Images were taken every 10s and play rate is 5fps.

**Movie S7: kinesin-1::3xsplitGFP (UNC-116::3xsplitGFP) are enriched at the leading end of mitochondria that transported anterogradely in the axon**

Lateral view of the synaptic region in the axon. Distal end is to the left. kinesin-1::3xsplitGFP in green and mito::TagRFP in magenta. A mitochondrion is moving anterogradely, with kinesin-1::3xsplitGFP at the leading end for most of the time. Note the presence of cytoplasmic pool of kinesin-1::3xsplitGFP. Images were taken every 10s and play rate is 5fps.

**Movie S8: RIC-7::7xsplitGFP are associated with mitochondria that enter the axon in *mtx-2* mutant, but is not enriched at the leading end**

Ventral view of the cell body with the axon to the right. The cell body is out-of-focus, which is indicated by the asterisk. RIC-7::7xsplitGFP in green and mito::TagRFP in magenta. RIC-7::7xsplitGFP is associated with the mitochondrion that moves anterogradely, but no longer accumulates at the leading end. Images were taken every 10s and play rate is 5fps.

**Movie S9 3D structure comparison by Dali.**

The structure of RIC-7b predicted by Alphafold was searched against the human proteome on the Dali server (http://ekhidna2.biocenter.helsinki.fi/dali/). The top two hits were OtulinL and Otulin. 3D structure alignment was shown in this movie, with RIC-7b in green and Otulin in orange. The predicted 3D structure of RIC-7b C-term is highly similar to that of Otulin.

